# An inosine triphosphate pyrophosphatase safeguards plant nucleic acids from aberrant purine nucleotides

**DOI:** 10.1101/2022.02.24.481826

**Authors:** Henryk Straube, Jannis Straube, Jannis Rinne, Markus Niehaus, Claus-Peter Witte, Marco Herde

## Abstract

- In plants, inosine is enzymatically introduced in some tRNAs but not in other RNAs or DNA. Nonetheless, our data show that RNA and DNA from *Arabidopsis thaliana* contain (deoxy)inosine, probably derived from non-enzymatic adenosine deamination in nucleic acids and usage of (deoxy)inosine triphosphate (dITP and ITP) during nucleic acid synthesis.
- We combined biochemical approaches, sample preparation and LC-MS, as well as RNA-Seq to characterize a plant INOSINE TRIPHOSPHATE PYROPHOSPHATASE (ITPA) from *Arabidopsis thaliana*, which is conserved in many organisms, and investigated the sources of deaminated purine nucleotides in plants.
- ITPA dephosphorylates deaminated nucleoside di- and triphosphates to the respective monophosphates. *ITPA* loss-of-function causes inosine di- and triphosphate accumulation *in vivo* and an elevated (deoxy)inosine content in DNA and RNA, as well as salicylic acid (SA) accumulation, early senescence and upregulation of transcripts associated with immunity and senescence. Cadmium-induced oxidative stress leads to more ITP in the wildtype, and this effect is enhanced in *itpa* mutants, suggesting that ITP originates from ATP deamination.
- ITPA is part of a molecular protection system, preventing accumulation of (d)ITP, its usage for nucleic acid synthesis, and probably nucleic acid stress leading to SA accumulation, stress gene induction and early senescence.

## INTRODUCTION

Spontaneous or enzymatically catalyzed reactions can lead to damaged or aberrant metabolites (Lerma-Ortiz et al., 2016; Crécy-Lagard et al., 2018; Carole L Linster et al., 2013). In recent years, several advances have been made in the field of damaged plant metabolites (Hanson et al., 2016), like the discovery of enzymes repairing NADH / NADPH hydrates (Niehaus et al., 2014), removing damaged glutathione (Niehaus et al., 2019) and oxidized deoxyguanosine triphosphate (8-oxo dGTP; Dobrzanska et al., 2002; Yoshimura et al., 2007). One of the most prominent metabolite classes that are prone to become damaged are nucleotides (Michael Y. Galperin et al., 2006; Nagy et al., 2014; Rudd et al., 2016; Rampazzo et al., 2010). In non-plant organisms, several enzymes have been identified that repair or remove aberrant and often toxic nucleotides, like DEOXYURIDINE TRIPHOSPHATE PYROPHOSPHATASE (DUT1), INOSINE TRIPHOSPHATE PYROPHOSPHATASE (ITPA), 8-OXO DEOXYGUANOSINE TRIPHOSPHATE PYROPHOSPHATASE (8-oxo dGTPase) and MULTICOPY ASSOCIATED FILAMENTATION (MAF) family proteins (Nagy et al., 2014). However, little is known about such enzymes in plants. While plant homologues of DUT1 and 8-oxo dGTPase (Dubois et al., 2011; Yoshimura et al., 2007) have been characterized, a plant ITPA has not been described.

ITPA enzymes from *Escherichia coli* and *Homo sapiens* remove deaminated purine nucleotide triphosphates (Lin et al., 2001; Davies et al., 2012). They exclusively hydrolyze the phosphoanhydride bond between the alpha- and beta-phosphate of (deoxy)inosine triphosphate (ITP and dITP) probably derived from deamination of (deoxy)adenosine triphosphate (ATP and dATP) or from spurious phosphorylation of inosine monophosphate (IMP). Xanthosine triphosphate (XTP), presumably derived from deamination of guanosine triphosphate (GTP) or spurious phosphorylation of xanthosine monophosphate (XMP), is also a substrate. ITPA generates the corresponding nucleotide monophosphate and pyrophosphate. Whereas (d)ITP and XTP are aberrant nucleotides, IMP and XMP are canonical intermediates of guanosine monophosphate (GMP) synthesis in most organisms and in plants XMP is additionally a starting point of purine nucleotide catabolism (Heinemann et al., 2021).

Deamination of cytosine to uracil is well known to occur in nucleic acids (Lindahl and Nyberg, 1974; Frederico et al., 1990) but purine bases can also be deaminated. *In vitro*, the deamination of adenine to hypoxanthine in DNA occurs at 2 to 3% of the rate observed for cytosine to uridine conversion (Lindahl, 1993). The deamination of free nucleotides is less well documented. Purine deamination was shown to occur not only in the nucleic acids but also in the pool of free nucleotides under heat stress (Karran and Lindahl, 1980). In general abiotic stresses, like salinity, water stress, extreme temperature, wounding, UV radiation or the exposure to heavy metals (Corpas et al., 2011; Valderrama et al., 2007) may influence the rate of nucleotide deamination. These stresses generate reactive nitrogen or oxygen species (RNS, ROS) (Corpas et al., 2011; Valderrama et al., 2007) which are mainly discussed in the context of protein modification and lipid peroxidation but are also likely damaging other metabolites although this has rarely been shown so far (Demidchik, 2015).

ITP and dITP are substrates of RNA and DNA polymerases, respectively. Random inosine or deoxyinosine in nucleic acids can thus not only be derived from deamination of adenosine or deoxyadenosine in the polymer but may also be incorporated during the polymerization. When DNA, containing deaminated sites, is replicated, mutations may be introduced. Deoxycytidine can be incorporated opposite of deoxyinosine and often thymidine is placed opposite of deoxyxanthosine (Kamiya, 2003). This misincorportion is not directly detrimental for *E. coli* (Budke and Kuzminov, 2006) because it is rapidly repaired. However, upon stimulation of deamination, the constant deployment of the repair machinery leads to strand breaks and chromosomal rearrangements (Budke and Kuzminov, 2010). By contrast, deoxyinosine in human DNA causes mutations and genome instability even when deamination is not stimulated (DeVito et al., 2017; Yoneshima et al., 2016). During RNA synthesis in non-plant organisms, incorporation of ITP has been shown to significantly reduce transcriptional velocity. Inosine in RNA alters RNA structure and stability and leads to mistranslation (Thomas et al., 1998; Ji et al., 2017). Apart from effects on nucleic acids, ITP and XTP can interfere with various cellular processes like G-protein signaling, microtubule assembly and ATP-dependent metabolic reactions in mammals and plants (Muraoka et al., 1999; Klinker and Seifert, 1997; Carnelli et al., 1992). Furthermore, ITP inhibits the TOPOISOMERASE II in *Drosophila melanogaster* (Osheroff et al., 1983), and interacts with the F1-subunit of the ATP synthase from *E. coli* (Weber and Senior, 2001).

ITP has been detected *in vivo* in only a few occasions. Human erythrocytes accumulated radioactive ITP after incubation with [^14^C] adenine, [^14^C] guanine or [^14^C] hypoxanthine, indicating that a biosynthetic route to deaminated purine nucleotide triphosphates might exist (Fraser et al., 1975). Interestingly, a loss of ITPA function in erythrocytes of mice leads to a substantial accumulation of ITP of up to 10 to 25% of the ATP pool (Behmanesh et al., 2009). However, erythrocytes are a special cell type which lack nuclear DNA and seem to have a particular nucleotide metabolism. It is technically challenging to detect ITP and XTP at all. In human cancer cells, both were detected with a highly sensitive method in the fmol/mg protein range. Here, ITP was around 5 to 10 times more abundant than XTP (Jiang et al., 2018). To date, the corresponding deoxyribonucleotide triphosphates, dITP and deoxyxanthosine triphosphate (dXTP), have never been detected in any organism.

Although these metabolites are low abundant, they can be incorporated into nucleic acids. The frequency of inosine in RNA is about one in 10^5^ nucleosides in wild type *E. coli*, while deoxyinosine in DNA is 10-fold less abundant (Pang et al., 2012). In mutants of rdgB, the bacterial ITPA homolog, the inosine content of RNA increased approximately 10-fold and the deoxyinosine of DNA rose about 5-fold compared to the wild type (Pang et al., 2012). In *ITPA* knockout-mice, the inosine to adenosine ratio was about one per 100 nucleosides in RNA (Behmanesh et al., 2009). In mammals, ITPA deficiency causes drastic phenotypes. In humans it leads to a lethal Martsolf-like syndrome with neurological and developmental abnormalities (Handley et al., 2019). Mice without a functional ITPA had RNA with an elevated inosine content and died shortly after birth (Behmanesh et al., 2009). However, bacteria lacking ITPA are viable (Burgis et al., 2003). Recently, a homolog of ITPA in *Cassava brown streak virus*, a plant-infecting RNA virus, was demonstrated to be an important factor during viral infection (Tomlinson et al., 2019).

In this study we identified ITPA of *A. thaliana* and biochemically characterized the enzyme, showing that it catalyzes the dephosphorylation of deaminated purine nucleotides. We analyzed mutants of *ITPA* demonstrating that the enzyme safeguards nucleic acids from incorporation of (d)ITP. We were able to detect ITP and IDP accumulation in the mutants and investigated the metabolic source of ITP and IDP. In contrast to *ITPA* mutants in mammals, the Arabidopsis mutants have only minor phenotypic alterations. They show slightly earlier senescence and have elevated concentrations of the defense phytohormone salicylic acid (SA), triggering a transcriptional response of genes associated with SA, systemic acquired resistance and aging.

## MATERIAL AND METHODS

### Plant cultivation

*Arabidopsis thaliana* and *Nicotiana benthamiana* plants were cultivated as described recently (Witte et al., 2005; Straube et al. 2021).

A segregating T-DNA insertion line from the SALK collection (*itpa-*1, SALK_052023) (Alonso et al., 2003) was acquired from the Nottingham Arabidopsis Stock Centre. A homozygous knockout line and a corresponding wild type were selected from the segregating population (see supporting methods).

A second knockout line was obtained via genome editing (*itpa-*2) according to Rinne et al. (2021, see supporting methods).

For root-length and germination assays, plants were grown in petri dishes on previously described medium (Straube et al., 2021; Niehaus et al., 2020).

Plants for the cadmium sulfate feeding experiment were grown hydroponically according to Batista-Silva et al. (2019), with small modifications. Plants were grown in viscous liquid medium (Straube et al., 2021; Niehaus et al., 2020) containing 0.4 % agar under long-day conditions. Cadmium treatment was performed for 24h by adding a 500 mM cadmium sulfate solution to a final concentration of 10 µm to the medium, respectively. This concentration was chosen according to Di Sanità Toppi and Gabbrielli (1999).

### DNA extraction, RNA extraction, genotyping and cloning

DNA for genotyping was isolated according to the BOMB-protocol using the TNA extraction from plants with the TNES/GITC lysis protocol (Oberacker et al., 2019). Magnetic beads were silica-coated beads, synthesized using the BOMB protocol for coating ferrite magnetic nano particles with silica oxide (Oberacker et al., 2019).

Total RNA was isolated using the Plant NucleoSpin II kit (Macherey-Nagel) according to manufacturer’s specifications.

Primers used in this study can be found in Table S1.

### Protein purification, quantification and enzymatic assay

C-terminal strep-tagged ITPA, AMPK3 and 4 (Chen et al., 2018) were transiently overexpressed in *N. benthamiana* and affinity purified after five days as described before (Werner et al., 2008). The protein concentration was determined by an in-gel quantification assay using a Li-cor Odyssey FC with Image Studio software. SDS gel electrophoresis and staining were performed as described previously (Witte et al., 2004).

Enzymatic assays for ITPA were performed using the EnzChek™ Pyrophosphate Assay Kit (ThermoFisher scientific) according to manufacturer’s instructions, with the exception of reducing the volume of all solution to one fifth of the instructions using High Precision Cells (Hellma Analytics).

AMPK3 and 4 activity was determined using a coupled enzyme assay with pyruvate kinase and lactate dehydrogenase (LDH, Sigma, P0294), measuring the production of NAD^+^ that results in absorption-decrease at 340 nm. For details see supporting methods.

### Subcellular localization

A construct encoding ITPA with C-terminal YELLOW FLUORESCENT PROTEIN (YFP) tag was transiently coexpressed in *Nicotiana benthamiana* with a construct encoding cytosolic β-ureidopropionase C-terminally fused to CYAN FLUORESCENT PROTEIN (CFP). Whole leaves were immersed in octadecafluor-decahydronaphthalin (Sigma-Aldrich, Littlejohn et al., 2010). The samples were analyzed using a Leica True Confocal Scanner SP8 microscope equipped with an HC PL APO CS2 40 × 1.10 water immersion objective (Leica, Germany) as described in Dahncke and Witte (2013). The acquired images were processed using Leica Application Suite Advanced Fluorescence software (Leica Microsystems). The immuno blot to analyze the construct was developed using a GREEN FLUORESCENT PROTEIN (GFP) -specific antibody. Co-localization was analyzed using the JACoP-plugin in ImageJ (Bolte and Cordelières, 2006).

### Metabolite analysis

The analysis of nucleotide concentration was performed as recently described with slight modifications (Straube et al., 2021), chromatographic parameters for the Hypercarb chromatography were changed as follows: flowrate, 0.6 ml min^-1^; column temperature, 35°C; injection volume, 20 µl. Gradient: 0 min, 100% A; 3 min, 100% A; 18 min, 70% A; 19 min, 0% A; 22 min, 0% A; 22.5 min, 100% A until 30 min. Transitions (precursor ions and product ions), fragmentor and collision energies employed for dITP, IDP, ITP, XMP and XTP are shown in Table 1.

**Table 1:** Precursor ions, transitions (quantifier, qualifier), fragmentor energy and collision energy for dITP, IDP, ITP, XMP and XTP

|  | Precursor Ion<br>[M+H]<sup>+</sup><br>(m/z) | Product Ion<br>(m/z) | Fragmentor | Collision Energy<br>(V) |
| --- | --- | --- | --- | --- |
| dITP | 492.99 | 217 | 115 | 10 |
|  |  | 137 |  | 30 |
|  |  | 81.1 |  | 30 |
| IDP | 429 | 137 | 80 | 10 |
|  |  | 97 |  | 22 |
| ITP | 508.99 | 137 | 120 | 30 |
|  |  | 97.1 |  | 30 |
| XMP | 365.05 | 212.9 | 84 | 3 |
|  |  | 153 |  | 15 |
|  |  | 97.1 |  | 17 |
| XTP | 525 | 244.3 | 17 | 40 |
|  |  | 97 |  | 40 |

The concentration of ATP and IMP was determined using the isotope dilution technique (Kuskovsky et al., 2019; Straube et al., 2021), while ADP, IDP, ITP and XMP were quantified by an external calibration curve in the respective plant matrix. Only calibration curves with an *R^2^-*value >0.99 were accepted. Calibration curves for IDP and ITP in buffer or matrix, as well as examples of wild type and knockout measurements can be found in the supplemental materials (Fig. S4). Salicylic acid was extracted and measured according to Cao et al., 2014 and Yang et al., 2017.

### Quantitative determination of (deoxy)inosine per (deoxy)adenosine level in total RNA and DNA

DNA was extracted using a modified CTAB-(hexadecyltrimethylammonium bromide) based protocol (see supporting methods). Total RNA was isolated using the Plant NucleoSpin II kit (Macherey-Nagel) according to manufacturer’s specifications.

Nucleic acids were digested using the Nucleoside Digestion Mix (NEB) with the following modifications: 5 μl Nucleoside Digestion Mix Reaction Buffer (10X) and 0.5 μl Nucleoside Digestion Mix were used, adding RNA orDNA and water depending on the respective concentration of the RNA or DNA up to a total volume of 50 μl. The mixture was incubated for 4 h at 37°C. A total of 1 μg RNA or 3 μg DNA was digested.

Digested RNA and DNA was measured by HPLC-MS as recently published (Traube et al., 2019), with minor modifications (Supporting methods).

### Analysis of chlorophyll content

Measurement of chlorophyll content was performed as described previously (Harris and Baulcombe, 2015).

### Cell death staining and quantification

Trypan blue staining was performed as published recently (Nuria Fernández-Bautista et al., 2016). To quantify the leaf area affected by cell death, RGB pictures of stained leaves were taken and stained cell areas were quantified using ImageJ.

### Sequence Analysis and Phylogenetic Analysis

ITPA sequences were identified using the Phytozome v12.1 webserver (https://phytozome.jgi.doe.gov/pz/portal.html) and the National Center for Biotechnology (NCBI) for all non-plant organisms employing BLASTP searches with the *Arabidopsis* amino acid sequence. The alignment and phylogenetic tree were constructed as described recently (Heinemann et al., 2021). The alignment used to construct the phylogenetic tree can be found in Fig. S1. For details, see supporting methods.

### RNAseq analysis

Total RNA from 35-days-old rosette leaves from *A. thaliana* Col-0 (‘wild type‘) and *itpa*-1 plants were send for Illumina mRNA sequencing to Novogene (Cambridge, United Kingdom). cDNA libraries were sequenced with 2×150 bp read length in *paired-end* mode. The total coverage of each sample is given in Table 2.

**Table 2:**
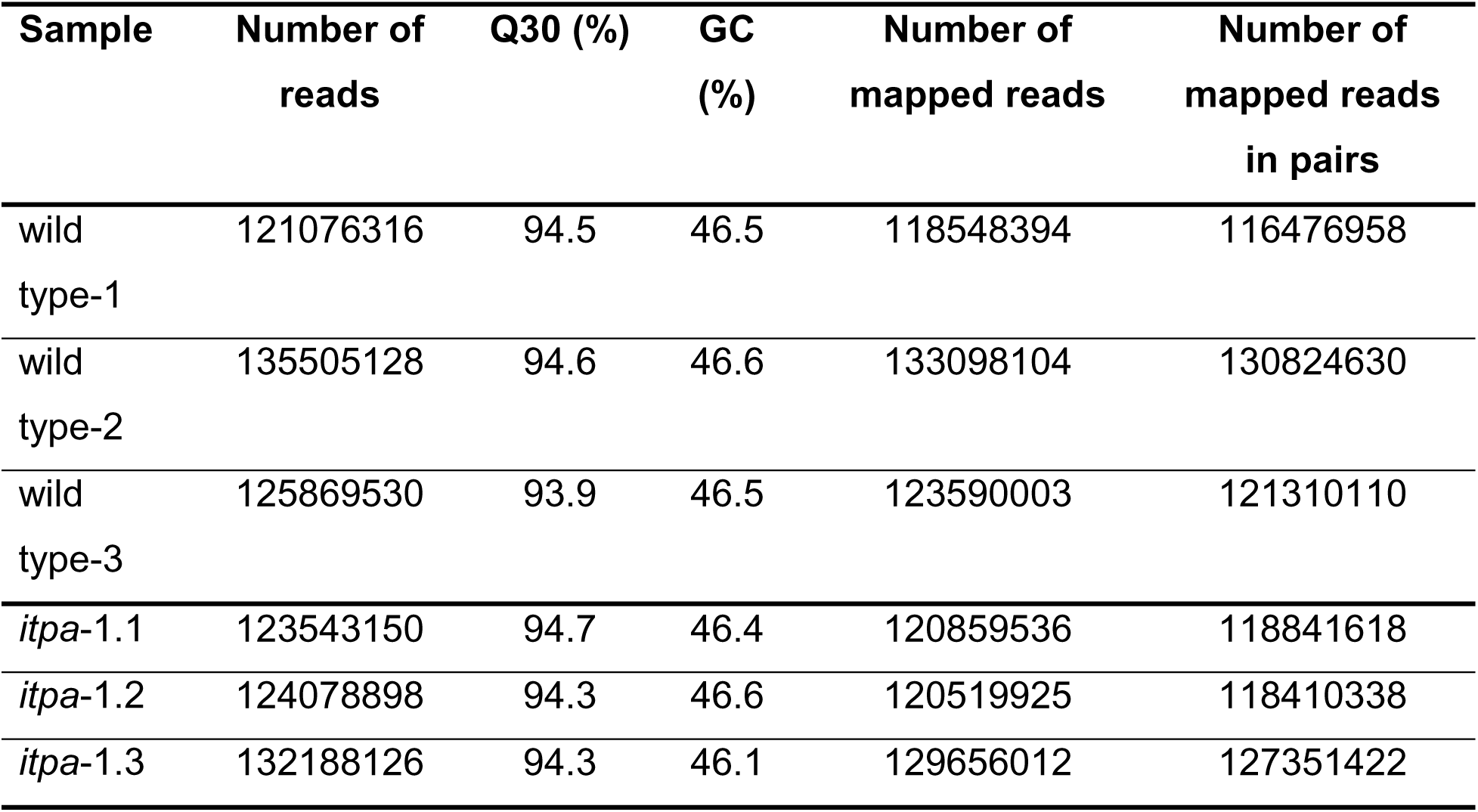
Sequencing and mapping results for RNAseq analysis

| Sample | Number of reads | Q30 (%) | GC (%) | Number of mapped reads | Number of mapped reads in pairs |
| --- | --- | --- | --- | --- | --- |
| wild type-1 | 121076316 | 94.5 | 46.5 | 118548394 | 116476958 |
| wild type-2 | 135505128 | 94.6 | 46.6 | 133098104 | 130824630 |
| wild type-3 | 125869530 | 93.9 | 46.5 | 123590003 | 121310110 |
| *itpa-1.1* | 123543150 | 94.7 | 46.4 | 120859536 | 118841618 |
| *itpa-1.2* | 124078898 | 94.3 | 46.6 | 120519925 | 118410338 |
| *itpa-1.3* | 132188126 | 94.3 | 46.1 | 129656012 | 127351422 |

Initial quality control of raw reads was performed with FastQC v0.11.9 (Andrews, 2010). After passing the quality control, reads were mapped to the *A*. *thaliana* (TAIR10) genome with the RNAseq-mapping algorithm included in the CLC Genomics Workbench v20.0.03 using default parameters. Genes with the following criteria were selected as differential expressed genes (DEG): log_2_ fold change (log_2_FC) ≥ 2, ≤ −2 (*itpa-1/*wild type); false discovery rate (FDR) ≤ 0.05 (Benajmin Hochberg correction). Log_2_ (reads per kilobase of model, RPKM) values were calculated as expression values. Single enrichment analysis (SEA) was done with the webtool agriGO v2.0 (Tian et al., 2017) for up- and down-regulated DEGs with the following parameters: Selected species: *Arabidopsis thaliana*; Reference: TAIR genome locus (TAIR10_2017); Statistical test method: Hypergeometric; Multi-test adjustment method: Hochberg (FDR); Significance level: 0.01; Minimum number of mapping entries: 10; Gene ontology type: Complete GO. A summary of DEGs and the results of the GO term enrichment analysis can be found in Table 2 and 3, respectively.

### Ribavirin treatment

The first fully developed true leaves of three-weeks-old *Nicotiana benthamiana* plants were fully inoculated with buffer (10 mM 2-(N-morpholino)ethanesulfonic acid, 10 mM MgCl_2_, pH 5.6) or buffer containing 500 μM ribavirin (1-[(2R,3R,4R,5R)-3,4-Dihydroxy-5-hydroxymethyl-oxolan-2-yl]-1,2,4-triazol-3-carboxamid). Samples were harvested after 2 h.

### Statistical analysis

Statistical analysis was performed as stated in Heinemann et al. (2021). The number of replicates, test values and multiplicity-adjusted p values are reported in Table S4.

## RESULTS

### Comprehensive phylogenetic analysis of inosine triphosphate pyrophosphatase in plants

We used the amino acid sequence of human ITPA to search with BLASTP for homologs in plants (Fig. S2). For *Arabidopsis thaliana,* the protein encoded at the gene locus At4g13720 was the only candidate with a high sequence similarity to the human ITPA. It has been suggested previously that this enzyme may represent a plant ITPA (Lin et al., 2001; James et al., 2021). A phylogenetic analysis revealed that ITPA is present in all kingdoms of life including plants and algae (Fig. 1, Fig. S1).

**Figure 1.**
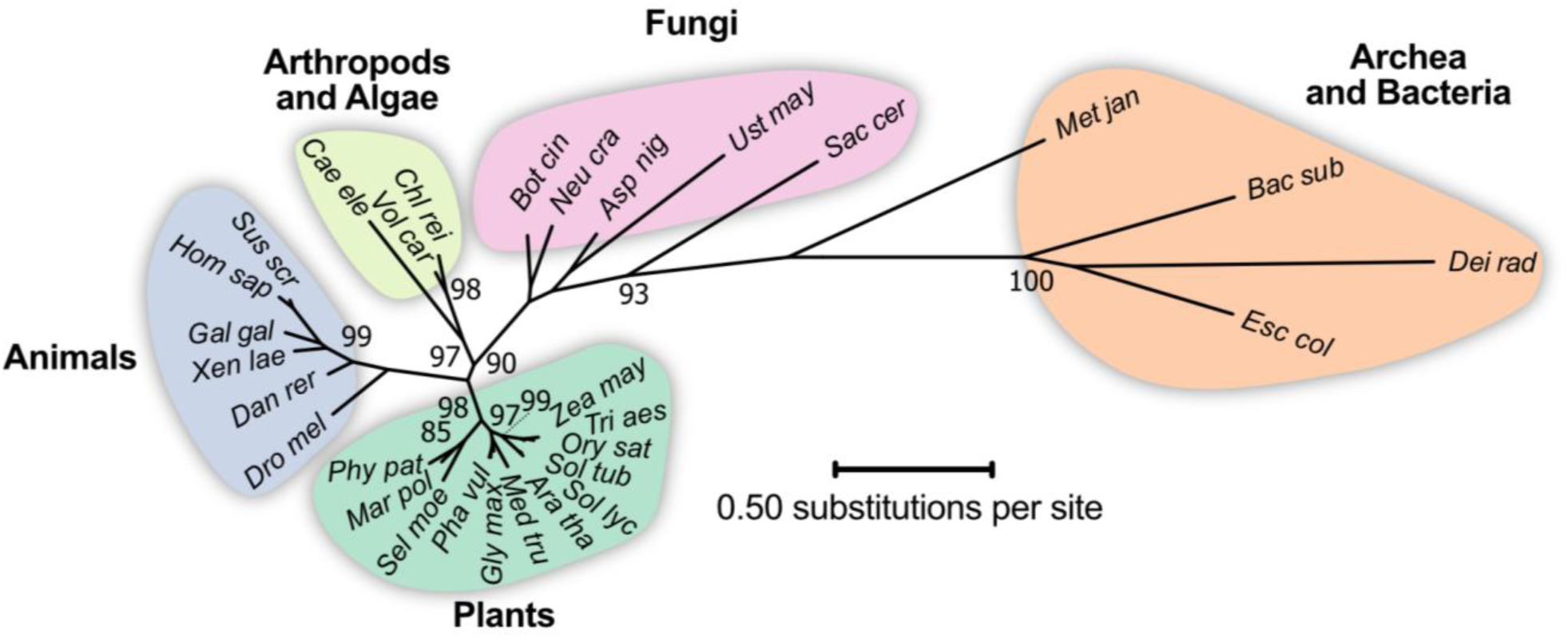
Phylogenetic analysis of ITPA in several organisms. An unrooted maximum likelihood tree was build using MEGAX, comprising model species from a wide taxonomic range. The tree with the best log-score is shown with branch length indicating the number of site-substitutions (scale-bar). Numbers at branches indicate the percentage of trees in which associated taxa clustered together in the bootstrap analysis (1000 bootstraps). Values below 80 are not shown. Species name abbreviations are listed in the legend of Fig. S1.

Several amino acid residues are highly conserved in ITPAs from different species. Many of these have been shown to be relevant for substrate binding, catalytic activity and dimerization (Fig. S1, S2; Stenmark et al., 2007; Gall et al., 2013). Nonetheless, there are amino acid residues that are only conserved in plant ITPAs (Fig. S1 and S2). Among these, K/R183 and K/R 189 can be acetylated in Arabidopsis (Liu et al., 2018), but the function of these posttranslational modifications is unclear.

### ITPA has atypical characteristics for an ITP pyrophosphatase

ITPAs are described in other organisms to hydrolyze the phosphoanhydride bond between the alpha and beta phosphate groups of inosine triphosphate resulting in inosine monophosphate (IMP) and pyrophosphate (Fig. 2a).

**Figure 2.**
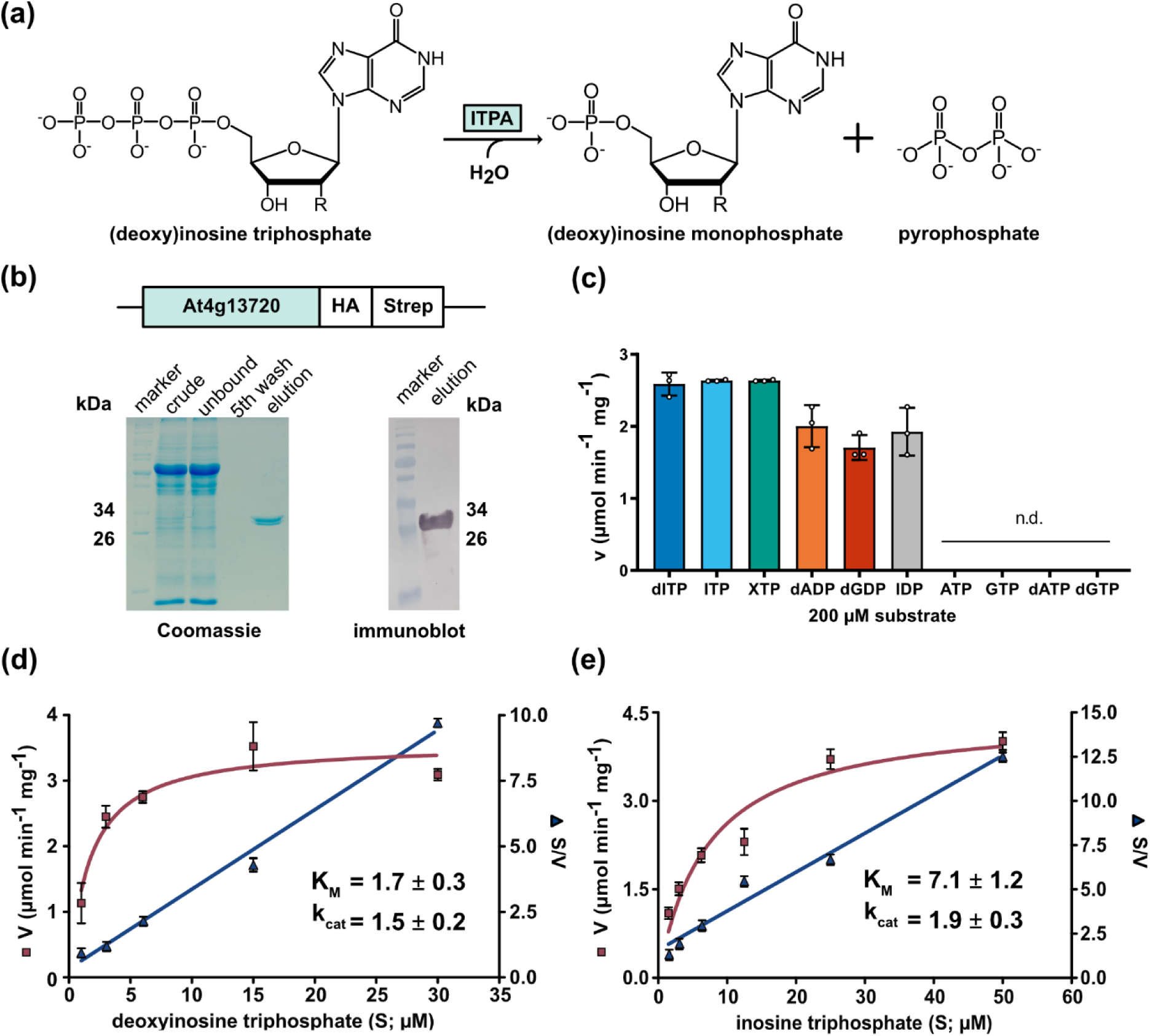
Characterization of the enzymatic properties of ITPA. (a) Reaction scheme of ITPA with ITP. (b) Affinity purified, C-terminal Strep-tagged ITPA on a Coomassie stained gel (left) and detection of ITPA by immunoblotting using an antibody against the Strep-tag (right). Numbers indicate the respective size in kilodalton (kDa). crude, centrifuged extract; unbound, centrifuged extract after incubation with Strep-tactin matrix; 5th wash, supernatant after the fifth wash of the affinity matrix; elution, eluate of the affinity matrix; boil off, supernatant after boiling the affinity matrix in SDS loading buffer. (c) Specific activity of ITPA with different substrate at a concentration of 200 µM. (d) Determination of kinetic constants for the ITPA catalyzed reaction of dITP to dIMP. Kinetic data was fitted using the Michaelis-Menten equation (orange) or by linear regression for the Hanes plot (S/V, purple). Error bars are SD, *n* = 3 reactions. S, substrate concentration; V, enzymatic activity. (e) The same as in (d), but using ITP as substrate.

To assess the activity of the plant enzyme, a C-terminal Strep-tagged variant of ITPA was transiently expressed in *Nicotiana benthamiana* and affinity purified (Fig. 2b). Interestingly, two forms of the protein were purified represented by two bands in a Coomassie-stained SDS-PAGE, possibly indicating a posttranslational modification. The identity of the purified protein was confirmed by immunoblot with an antibody against the Strep-tag. The specific activity of the purified enzyme was determined with several substrates at concentrations of 200 μM, including (deoxy)ribonucleotide di-, and triphosphates, *para*-nitrophenyl-phosphate and nicotinamide adenine dinucleotide (Fig. 2c, Table S5). An enzymatic activity could only be observed with dITP, ITP, inosine diphosphate (IDP) and xanthosine triphosphate (XTP), as well as with deoxyadenosine diphosphate (dADP) and deoxyguanosine diphosphate (dGDP). In contrast to the ITPA from *H. sapiens,* the ITPA of *A. thaliana* efficiently dephosphorylates IDP (Davies et al., 2012). For dITP and ITP the kinetic constants were determined (Fig. 2d, e). With ITP as substrate a K_M_ of 7.1 ± 1.2 μM and a turnover number of 1.9 ± 0.25 s^-1^ were measured. With dITP the K_M_ was about four-fold lower at 1.7 ± 0.3 μM while the turnover number was similar at 1.5 ± 0.17 s^-1^. Thus, Arabidopsis ITPA has catalytic efficiencies of 2.6 x 10^5^ mM^-1^ s^-1^ for ITP and 8.8 x 10^5^ mM^-1^ s^-1^ for dITP (Table 3).

**Table 3:**
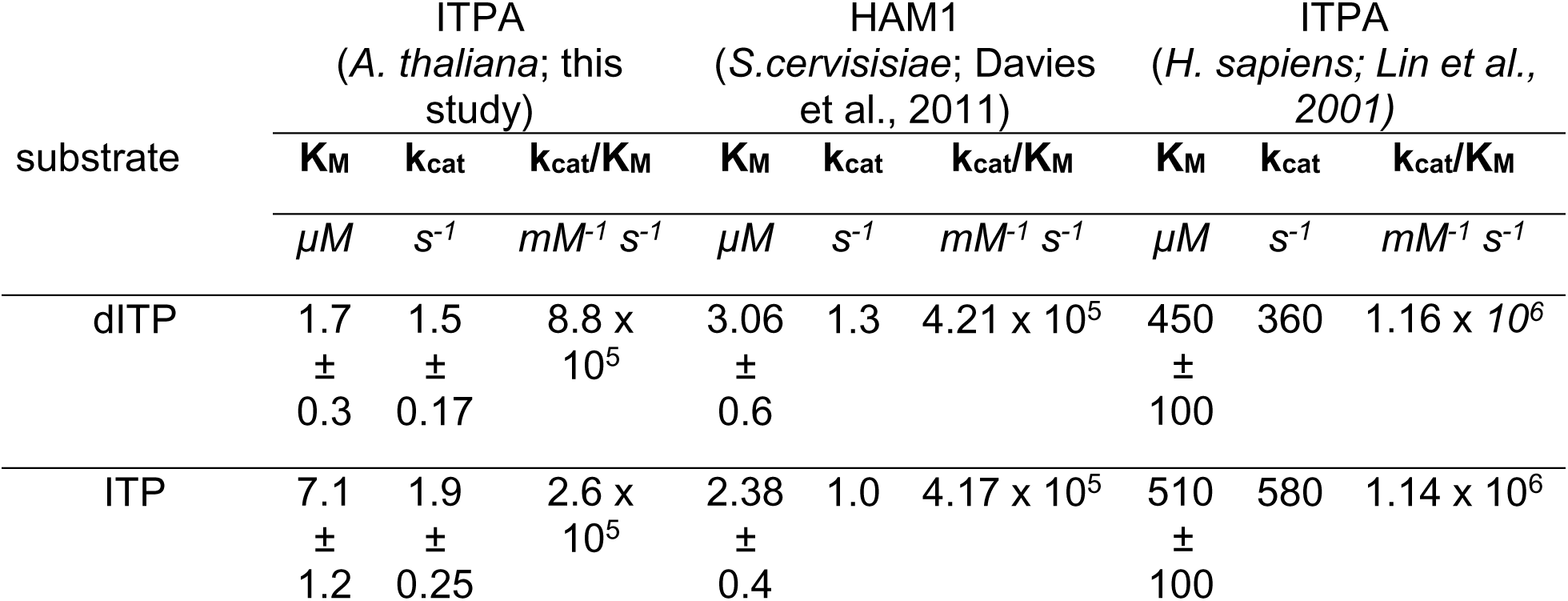
Comparison of kinetic parameters of plant, fungi and mammal ITPA enzymes

### ITPA mutants show an altered phenotype during senescence

To further investigate the role of ITPA *in vivo*, two mutant lines were created. One T-DNA line with an insertion in an exon of *ITPA* was available from the SALK collection (SALK_053023; Fig. 3a).

**Figure 3.**
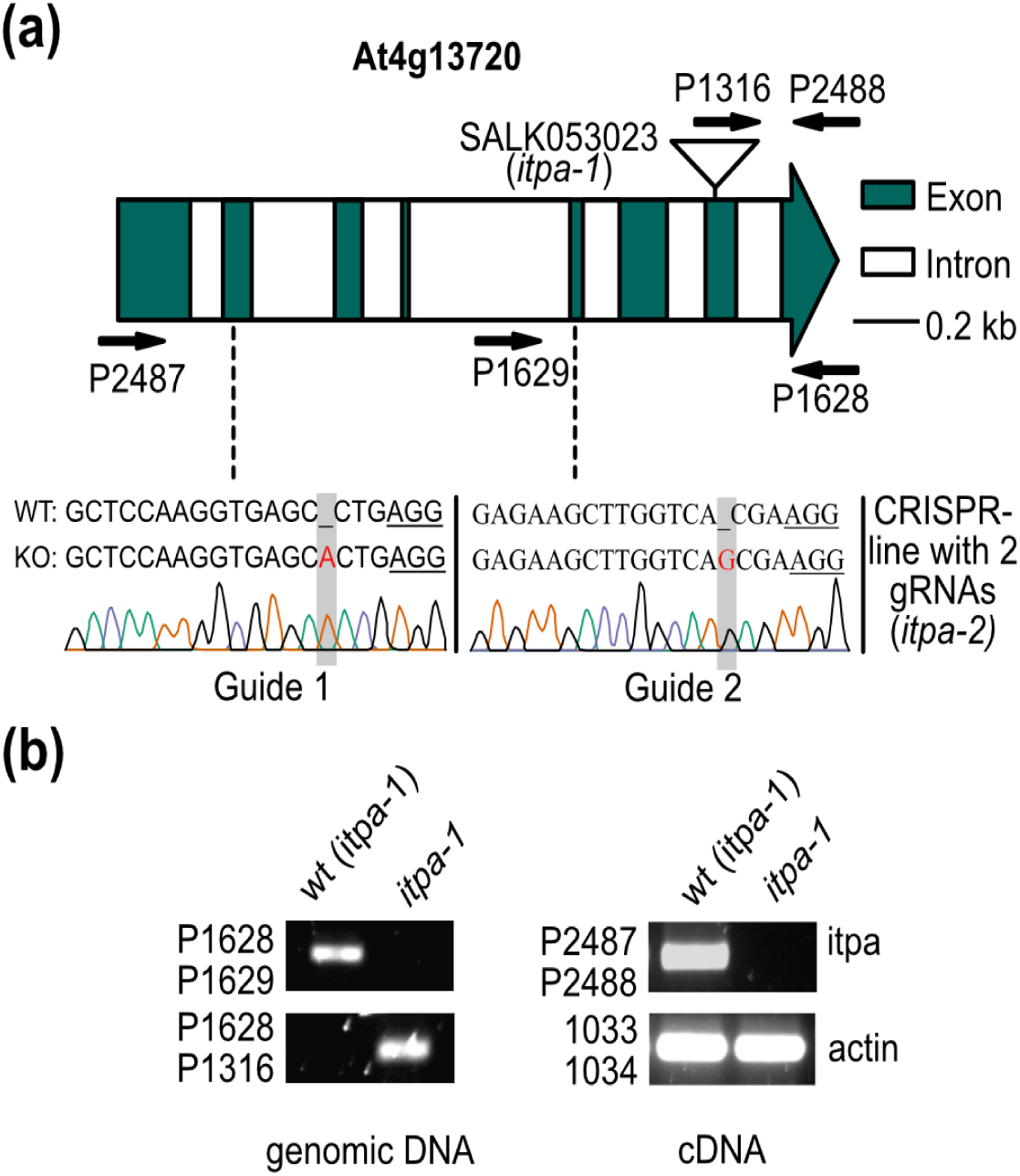
Characterization of *itpa* mutant lines. (a) Organization of the At4g13720 locus encoding ITPA and positions of T-DNA insertion sites (SALK053023; Δ *itpa-1*), as well as binding site of the guide RNAs (gRNAs) used to generate Δ *itpa-2*. The respective gRNAs are shown below as well as the single base pair insertions observed in *itpa-2*. Exons are shown as green blocks, introns as white blocks. Arrows indicate primer positions. (b) PCR-analysis of the wildtype and *itpa-1* alleles (left panel), as well as RT-PCRs to assess the presence of intact mRNA in homozygous *itpa-1* lines (right panel) and the corresponding wildtype selected from a segregating mutant population. The mRNA of the actin2 gene (At3g18780) was amplified as control.

We identified plants with a homozygous insertion of the T-DNA and observed that they lacked an intact transcript (Fig. 3b). This mutant was named *itpa-1*. A second line (*itpa-2*) was generated using CRISPR, resulting in a plant with two homozygous single base pair insertions causing frameshifts, one in exon 2 and one in exon 5. From the segregating population harboring the editing construct, *itpa-2* plants without the transgene were selected and used for further characterization. Both mutant lines had no obvious developmental phenotypes (Fig. S3 a-d) but showed an early onset of senescence at 40 days after germination (Fig. 4a) and a slightly increased senescence at an age of eight weeks with around 50% less chlorophyll in the oldest leaves, as well as an increased percentage of dead cells per leaf in *itpa* plants when compared to the wild type (Fig. 4b-d). The plant fresh weight, the number of seeds per silique and the length of the siliques were not different between the mutants and the wild type (Fig. S3 b-d). The expression of *ITPA* is rather constitutive like a typical housekeeping gene and does not correlate with age according to the eFP browser (Winter et al., 2007).

**Figure 4.**
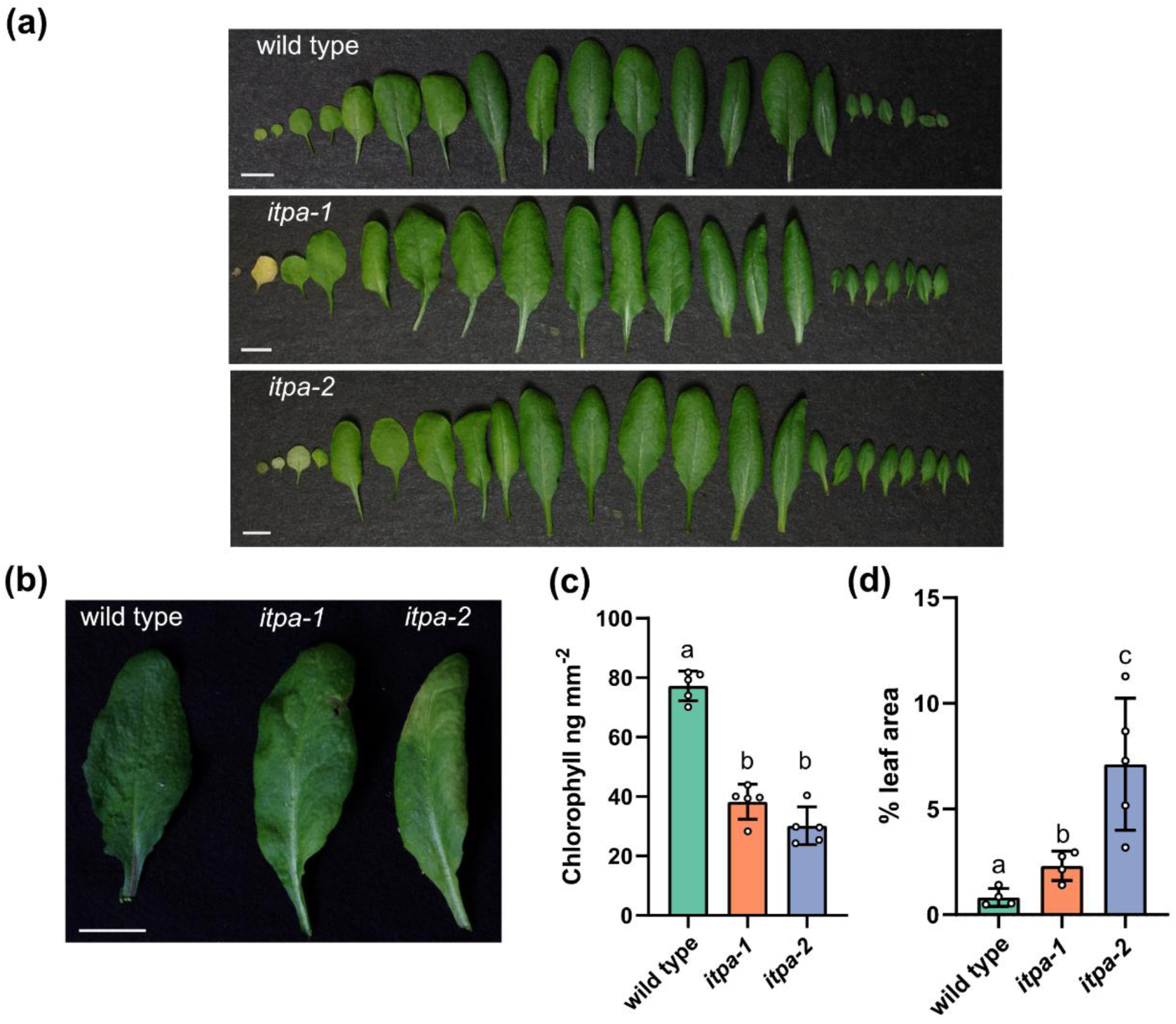
Phenotypic comparison of wild type, *itpa-1* and *itpa-2* plants. (a) Rosette leaves of 40-day-old wild type, *itpa-1* and *itpa-2* plants grown under long day conditions. Bars = 10 mm. (b) Representative picture of the oldest leaves from 56-day-old plants of different genotypes used in (c). (c) Chlorophyll content of the respective oldest leaves from 56-day-old plants of different genotypes. *n* = 5, every replicate represents a different plant of the same genotype. (d) Leaves of 56-days-old wild type, *itpa-1* and *itpa-2* plants were stained by trypan blue and dark pixels per leaf-area were measured using ImageJ. Error bars are SD, *n* = 4 - 5. Statistical analysis was performed by a two-sided Tukey’s pairwise comparison using the sandwich variance estimator. Different letters indicate *p* values < 0.05. All *p* values can be found in the Table S4.

### ITPA is localized in the cytosol, nucleus and putatively in plastids

To determine the subcellular localization of ITPA, cells of *N. benthamiana* co-expressing a C-terminally YFP-tagged variant of ITPA and cytosolic β-ureidopropionase C-terminally fused to CFP. ITPA was clearly localized in the cytosol and nucleus, co-localizing with the cytosolic marker as the signals correlate with an r-value of 0.98, additionally a good fit in the cross-correlation analysis was observed (Fig. 5a, b).

**Figure 5.**
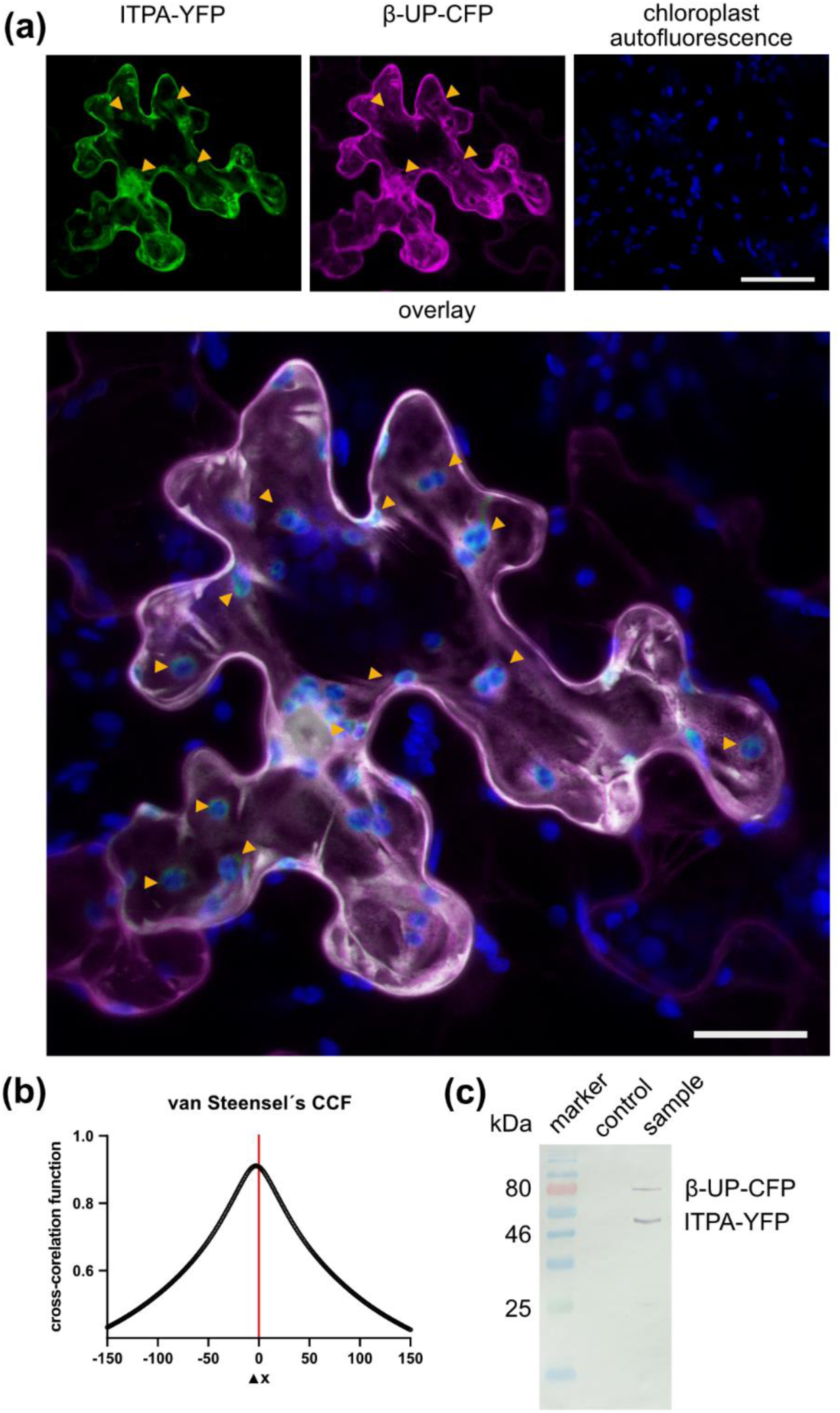
Subcellular localization of ITPA. (a) Confocal fluorescence microscopy images of leaf epidermal cells of N. benthamiana transiently coexpressing ITPA C-terminally fused to YFP (ITPA-YFP) and cytosolic β-ureidopropionase C-terminally fused to CFP (β-UP-CFP). Left panel, YFP fluorescence; middle panel, CFP fluorescence; right panel, chloroplast auto fluorescence. Bar = 50 µm. The large picture is an overlay of all three signals. Bar = 25 µm. Orange arrows indicate YFP signal not overlapping with the CFP signal. Analysis of the Pearsońs correlation coefficient gave an r-value of 0.98 for the YFP and CFP signal. The images are representative for at least 10 micrographs of different cells. (b) Van Steenseĺs cross-correlation analysis for the YFP and CFP signal. (c) Analyzing the stability of the ITPA-YFP and β-UP-CFP by an immunoblot developed with an GFP-specific antibody. The control was generated from infiltrated leaves with *A. tumefaciencs* carrying the RNA silencing suppressor P19.

Both fusion proteins are stable as shown by an immunoblot (Fig. 5c). Interestingly, while the plastids are completely dark in the channel corresponding to the CFP signal of the cytosolic marker, there is signal in the YFP channel at these locations, illustrated by the arrows. Furthermore, the YFP signal and the plastidic autofluorescense overlap, indicated by a cyan coloration in the overlay of micrographs (Fig. 5a). Although the signals do not coincide completely, the data indicate that ITPA might also be localized in the plastids. This is in line with the predictions of several bioinformatic analysis tools summarized in the SUBA4 database, predicting nuclear, cytosolic and plastidic localization for ITPA (Hooper et al., 2017). Additionally, ITPA was found in a proteomic analysis of the plastid proteome (Kong et al., 2011). A localization in mitochondria, also predicted by SUBA4, was not confirmed in our experiments.

### ITPA is essential for the catabolism of ITP and IDP *in vivo*

In 35-day-old plants, IDP and ITP were only detectable in *itpa-1* and *itpa-2* background but not in the wild type (Fig. 6a).

**Figure 6.**
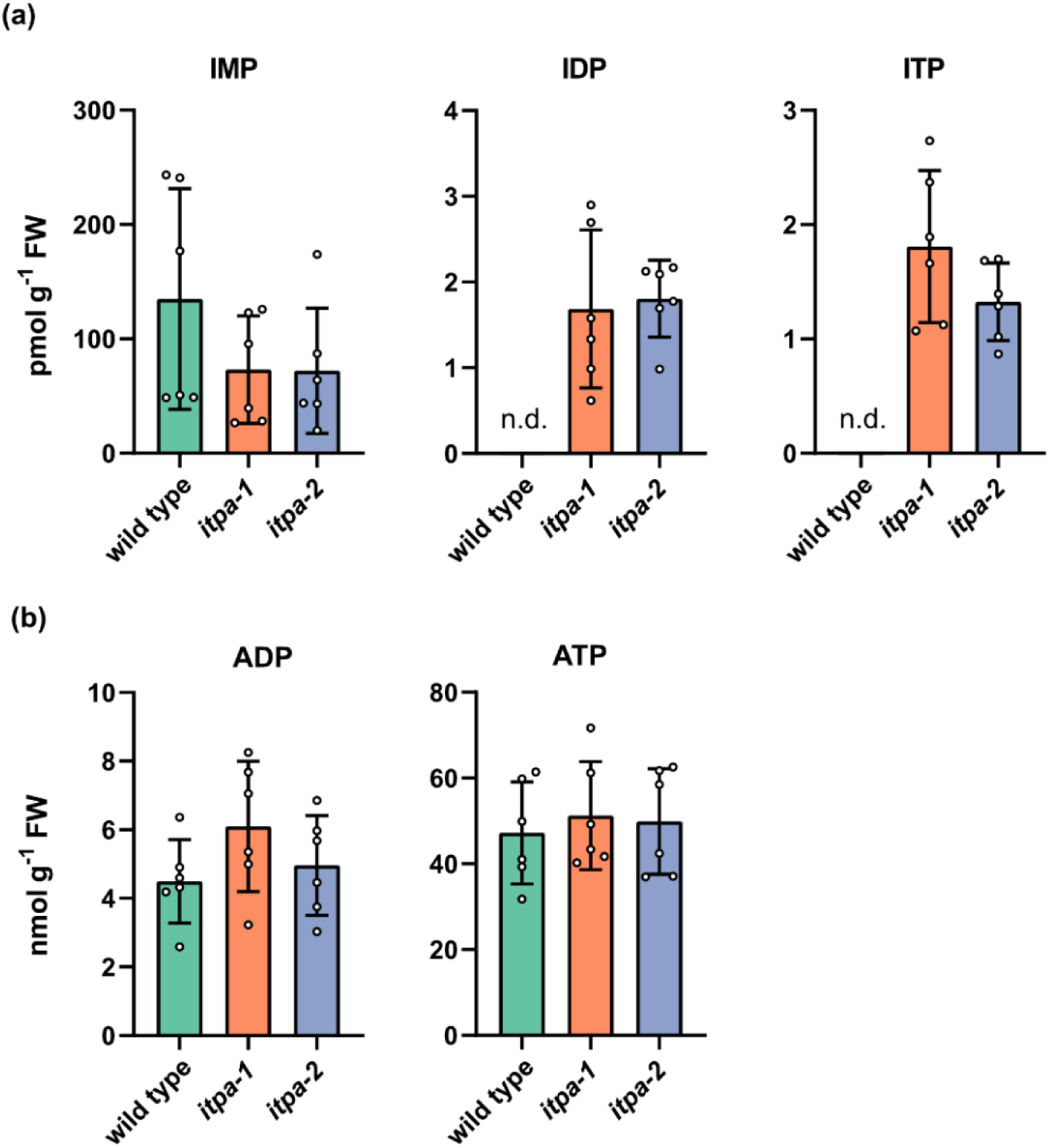
Loss of function of *itpa* causes an accumulation of IDP and ITP while adenylate concentrations remain unchanged. (a) Concentrations of inosylates in rosette leaves of 35-day-old wild type, *itpa-1* and *itpa-2* plants. (b) As in (A) but for ADP and ATP. Error bars are SD, *n* = 6, every replicate represents a different plant. Statistical analysis was performed by a two-sided Tukey’s pairwise comparison using the sandwich variance estimator. Different letters indicate *p* values < 0.05. All *p* values can be found in the Table S4. n.d. = not detected.

ITP and IDP were reliably measured because they were baseline separated from the isobaric ATP and ADP isotopes (Fig. S4). The *itpa-1* and *itpa-2* lines contained 1.7 and pmol g^-1^ IDP and 1.8 and 1.3 pmol g^-1^ ITP, respectively. This accounts for about 30 molecules of ITP per 10^6^ molecules of adenosine triphosphate (ATP) and between 300 and 400 molecules of IDP per 10^6^ molecules of adenosine diphosphate (ADP, Table 4).

**Table 4:** Ratios of ITP and IDP per million molecules of ATP and ADP, respectively

|  | *itpa-1* | *itpa-2* |
| --- | --- | --- |
| ITP/ 1 x 10<sup>6</sup> ATP | 36.6 ± 12 | 28,7 ± 11 |
| IDP/ 1 x 10<sup>6</sup> ADP | 300,8 ± 162 | 407,5 ± 178 |

The ITP concentration per cell was about 9.6 nM and that of IDP about 11.2 nM estimated according to Straube et al. (2021). Interestingly, the IMP concentration in both knockout lines appears to be lower in tendency although the data is statistically not significant (Fig. 6a). However, the difference in mean IMP concentrations between the wild type and *itpa* plants is by far greater than the sum of ITP and IDP accumulating in the mutants suggesting that ITP and IDP dephosphorylation is not a main contributor to the IMP pool. This is not surprising because IMP is a comparatively abundant intermediate of purine nucleotide biosynthesis and purine nucleotide catabolism (Witte and Herde, 2020; Baccolini and Witte, 2019). Although ITPA is active with dADP and dGDP *in vitro*, mutation of *ITPA* had no effect on the dADP and dGDP concentrations *in vivo* (Fig. S5a). The concentrations of other purine and pyrimidine nucleotides were also not altered in the mutant lines (Fig. 6B, Fig. S5a-c).

### What is the metabolic source of ITP and IDP?

Hitherto, the metabolic source of ITP and IDP is unknown. One possibility is that these metabolites arise from promiscuous enzyme activity (Hanson et al., 2016) of AMP kinases which in addition to AMP may also be able to phosphorylate IMP. Consistent with this hypothesis, it has been shown in *E. coli* and *Saccharomyces cerevisiae* that an increase of IMP content is concomitant with an accumulation of (deoxy)inosine in RNA and DNA (Pang et al., 2012). In plants, more IMP could also lead to more ITP and IDP, which would indicate that phosphorylation of IMP by kinases is the source of ITP and IDP. To test this hypothesis, we infiltrated leaves of *N. benthamiana* with ribavirin, an inhibitor of IMP DEHYDROGENASE (IMPDH, Keya et al., 2003). This enzyme represents a major sink of IMP in plant metabolism (Witte and Herde, 2020) and its inhibition may allow to manipulate the IMP concentration. *N. benthamiana* was chosen, because even in untreated wild type leaves ITP is measurable in this plant and thus concentration changes upon treatment can be monitored, whereas in Arabidopsis this metabolite is often only detectable in the *itpa* background (Fig. 6a). Ribavirin treatment for two hours caused a significant increase of the IMP content in leaves of *N. benthamiana* compared to the mock treatment with buffer, and a decreased concentration of downstream metabolites like XMP, GMP and in tendency GTP (Fig. 7 and Fig. S6).

**Figure 7.**
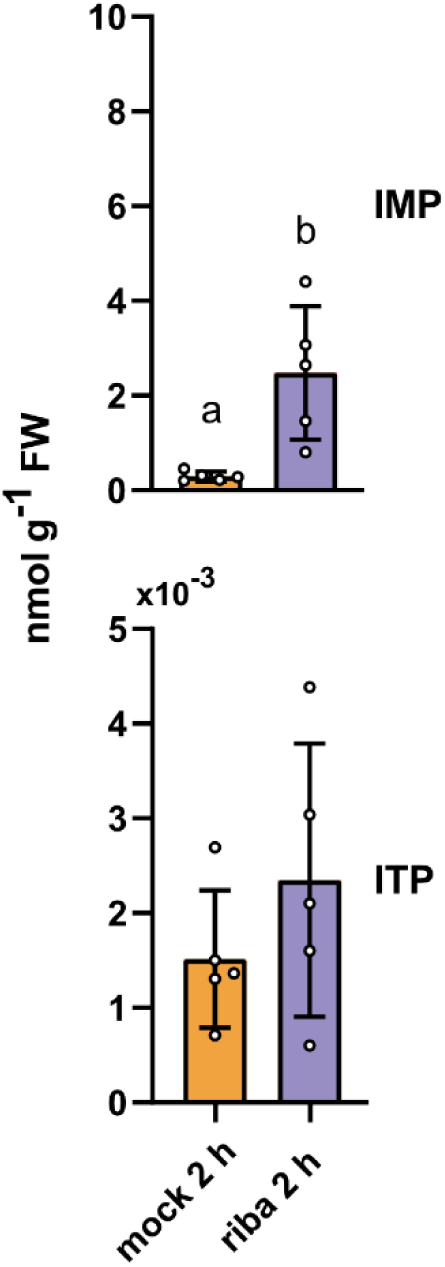
Manipulation of IMP content in leaves of *N. benthamiana* does not alter ITP concentrations. IMP (upper panel) and ITP (lower panel) in leaves of 21-day-old *N. benthamiana* plants. The leaves were either mock infiltrated with buffer on one leaf or infiltrated with buffer containing ribavirin (500 µM) on the opposite leaf. Error bars are SD, *n* = 5, every replicate representing a leaf from an independent plant. Statistical analysis was performed by a two-sided Tukey’s pairwise comparison using the sandwich variance estimator. Different letters indicate *p* values < 0.05. All *p* values can be found in the Table S4.

However, the ITP content did not change in any case, and this despite the fact that it was always several orders of magnitude lower than the IMP content. The data show that ITP levels are not directly dependent on IMP concentrations.

A biochemical approach was also used to investigate the question of possible aberrant IMP phosphorylation. We determined the enzymatic activities of the two cytosolic AMP kinases, AMPK3 (At5g50370) and AMPK4 (At5G63400) of *A. thaliana* with either 1 mM AMP or IMP as substrates (Lange et al., 2008; Chen et al., 2018; Fig. 8a, b). Although IMP is an *in vitro* substrate for both enzymes, the activity with IMP was only 3.9% of that with AMP for AMPK3 (Fig. 8a) and 5.2% for IMP versus AMP for AMPK4 (Fig. 8b).

**Figure 8.**
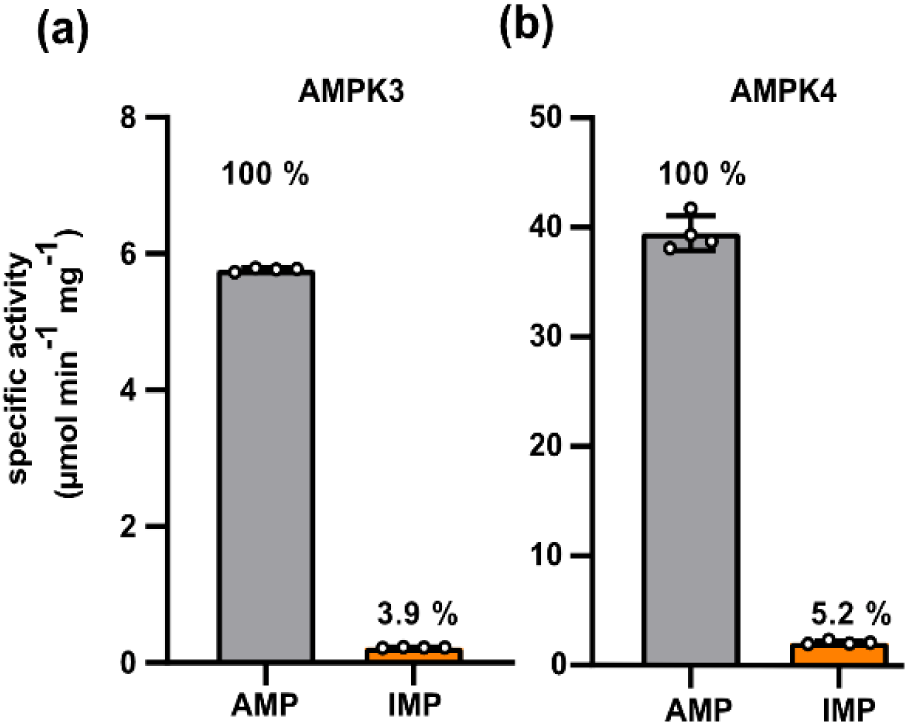
Enzymatic activity of AMPK 3 and 4 with AMP and IMP. Enzymes were transiently expressed in *N. benthamiana* as C-terminal Strep-tagged variants and affinity purified. (a) Specific activity of AMPK3 with 1 mM AMP or IMP in the presence of 1 mM ATP. Linear rates were observed in all cases. Error bars are SD. Each replicate represents an independent enzyme assay from the same enzyme preparation (*n* = 4). (b) Analogous data for AMPK4.

*In vivo* the relative activity with IMP will be lower because the cellular IMP pool is several orders of magnitude smaller than that of AMP. Nonetheless, the data show that AMPKs can phosphorylate IMP in principle but unlikely contribute strongly to the formation of ITP and IDP.

Aberrant metabolites can also be derived from non-enzymatic processes – e.g. from contact with reactive oxygen species (ROS; Hanson et al., 2016; Lerma-Ortiz et al., 2016). Heavy metals, such as cadmium, are known to cause ROS production in plants and comprise important anthropogenic pollutants (Di Sanità Toppi and Gabbrielli, 1999; Cho and Seo, 2005; Lin et al., 2007). It has been well established that cadmium-induced oxidative stress not only damages macro molecules such as the DNA directly, but also causes an increased rate of lipid peroxidation (Lin et al., 2007). To test whether this oxidative stress also has an impact on nucleotides, we grew wild type and *itpa-1* plants hydroponically for 14 days and applied cadmium sulfate (0 μM and 10 μM). Without Cd, wild type plants only accumulated small amounts of ITP, but with 10 μM Cd the ITP content rose significantly (Fig. 9). IDP was not detectable in the wild type. As expected, the mutant generally contained more IDP and ITP than the wild type and the application of Cd resulted in a further increase of these pools. The concentrations of ITP and IDP measured in this experiment in 14-day-old plants grown in a hydroponic system were comparable to those in 35-day-old plants grown on soil (Fig. 6a and Fig. 9). These data show ITP and IDP can be formed under stress conditions that involve ROS probably by deamination of ATP and ADP. Overall, our data suggest that the main source of ITP is not IMP phosphorylation but ATP deamination.

**Figure 9.**
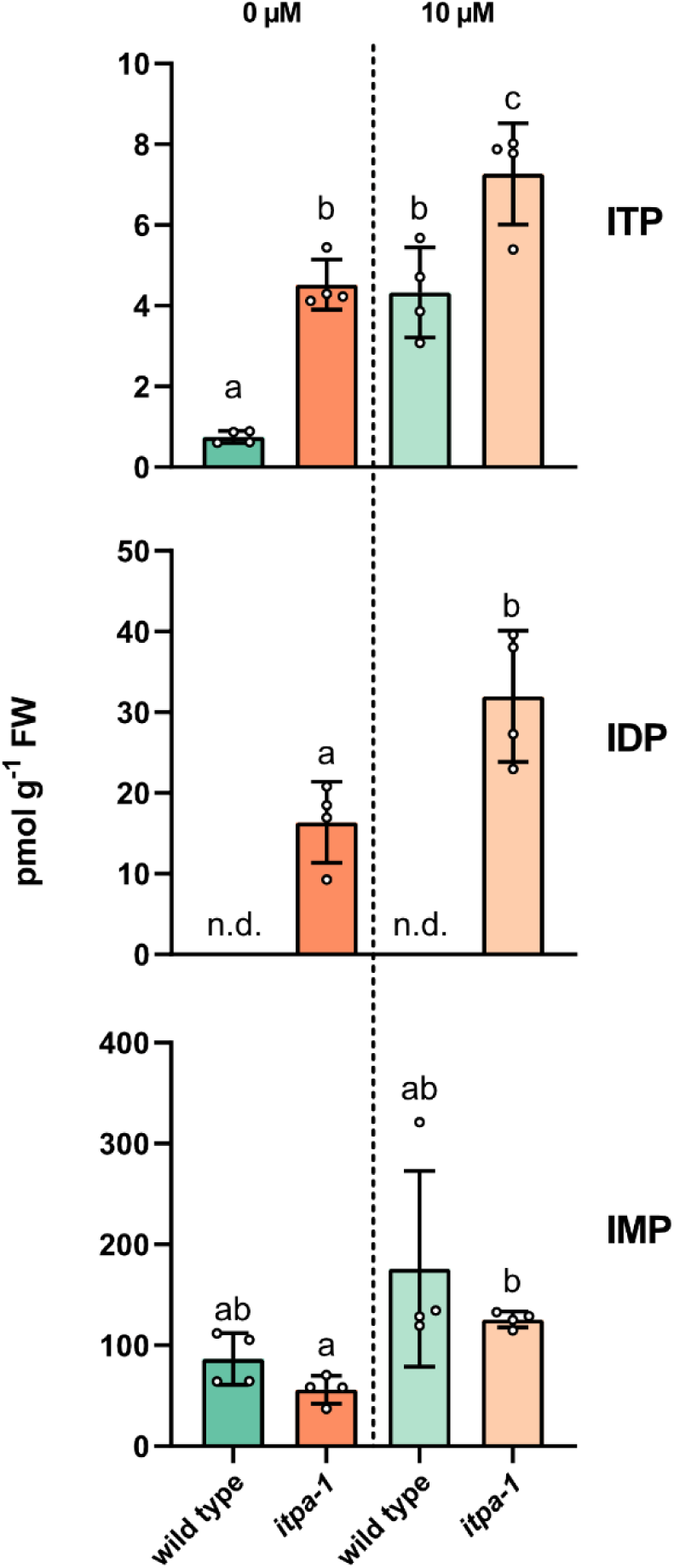
The effect of cadmium causing oxidative stress on inosylate concentrations. Mutant and wild type plants grown in a hydroponic system for 14 days were treated with 10 µM cadmium sulfate for 24 h or were left untreated. ITP, IDP and IMP were quantified. Error bars are SD, *n* = 4, every replicate represents a pool of plants from an independent hydroponic vessel. Statistical analysis was performed by a two-sided Tukey’s pairwise comparison using the sandwich variance estimator. Different letters indicate *p* values < 0.05. All *p* values can be found in the Table S4. n.d. = not detected.

### ITPA prevents (d)ITP incorporation into DNA and RNA

Measuring deoxyinosine (dI) in digested DNA of 7-day-old plants by liquid chromatography-mass spectrometry (LC-MS) revealed that both mutant lines accumulate dI in the DNA (Fig. 10a). Wild type DNA contained 290 ± 96 dI per million molecules deoxyadenosine (dA). This amount was more than doubled in the mutants (Table S6). In DNA from older plants (56 days) dI could not be detected in the wild type and accumulated in DNA of the mutants to approximately the same level as in seedlings. Therefore, a marked increase of dI accumulation in DNA with age could not be observed. Interestingly, the higher dI concentration in DNA of *itpa* plants demonstrates that a dITP pool likely exists in plants. Deoxyxanthosine was not detected in DNA.

**Figure 10.**
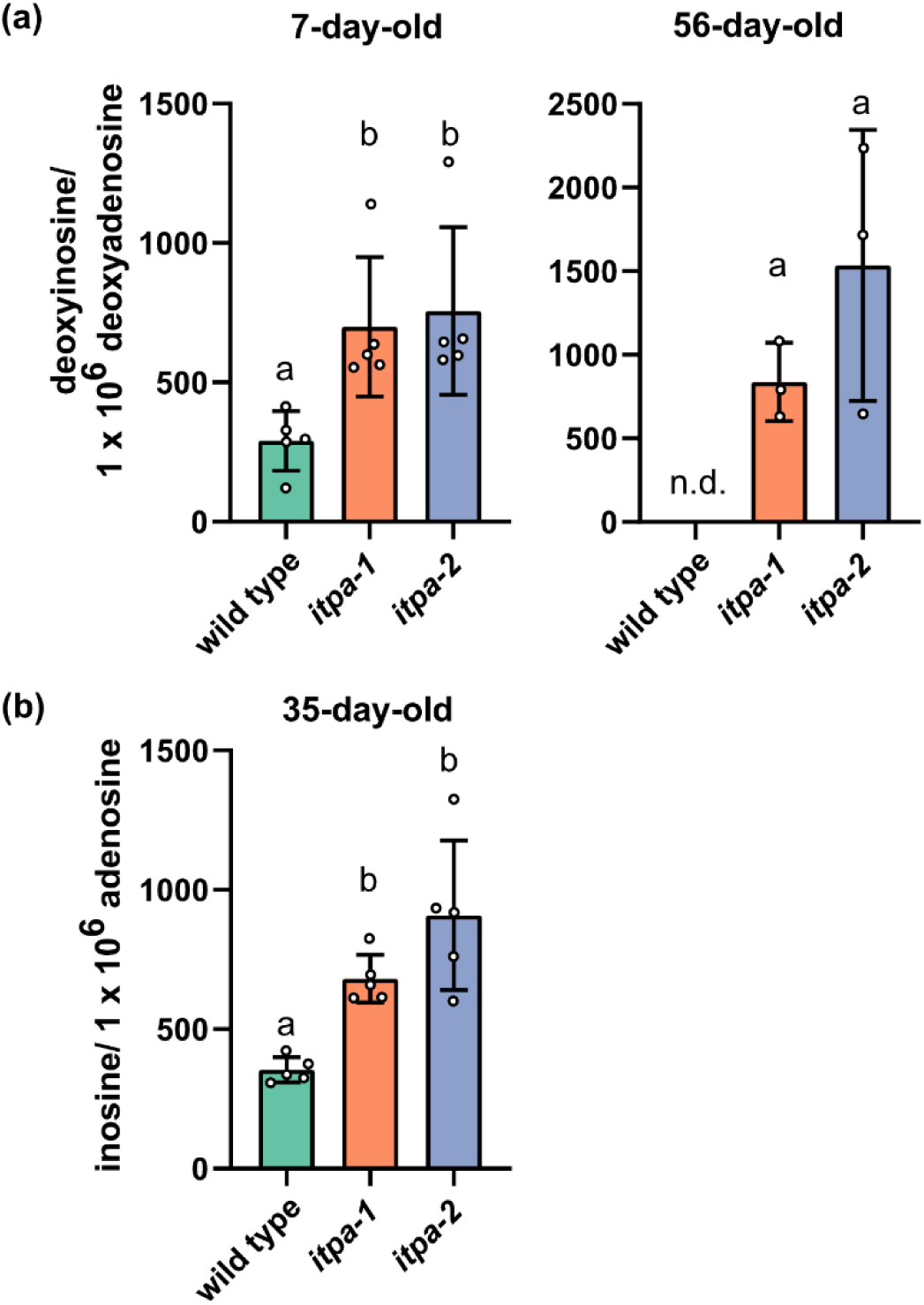
Concentration of deoxyinosine and inosine in nucleic acids of different genotypes. (a) Molecules of deoxyinosine per million molecules of deoxyadenosine in DNA isolated from pools of 7-day-old Arabidopsis seedlings grown on agar plates and 56-day-old Arabidopsis rosettes. Error bars are SD, *n* = 5 and 3. (b) Molecules of inosine per million molecules adenosine in total RNA isolated from 35-day-old Arabidopsis rosettes. Error bars are SD, *n* = 5. Every replicate represents a different plant. Statistical analysis was performed by a two-sided Tukey’s pairwise comparison using the sandwich variance estimator. Different letters indicate *p* values < 0.05. All *p* values can be found in the Table S4. n.d. = not detected.

In digested total RNA of 35-day-old plants, 405 ± 117 molecules of inosine per million molecules of adenosine were detected (Fig. 10b). The *itpa* lines contained about twofold more inosine in the RNA (Table S6) showing that also the RNA is protected by ITPA. In a pilot experiment, the RNA of younger, 21-day-old plants was analyzed and contained only 200 molecules inosine per million molecules of adenosine (Fig. S7), which is about half the amount found in 35-day-old plants. A similar tendency was observed for the *itpa-1* line. In contrast to deoxyinosine in DNA, it appears that the inosine content of RNA increases with age. Interestingly, inosine containing RNA can be specifically recognized and cleaved by an endonuclease in Arabidopsis, which may limit the amount of inosine that can accumulate in RNA (Endo et al., 2021). Xanthosine was not detected in RNA. The abundance of known damaged nucleosides like 8-oxo guanosine in RNA was not different between wild type and mutant plants. In summary the data demonstrate that ITPA is an enzyme required for the removal of dITP and ITP to protect DNA and RNA from random incorporation of (deoxy)inosine (Fig. 11).

**Figure 11.**
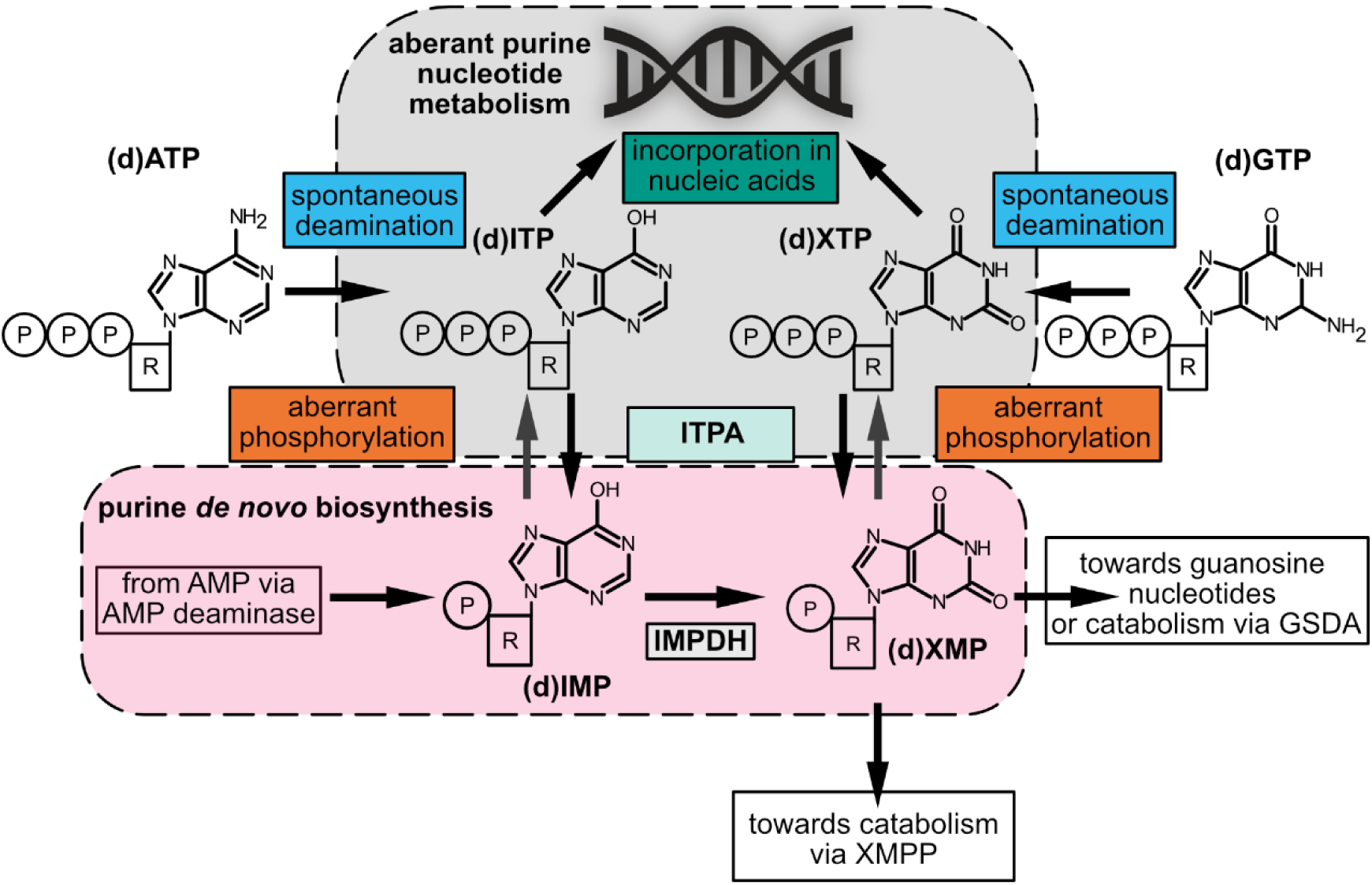
Overview of deaminated purine nucleotide metabolism and its interplay with purine *de novo* synthesis. Enzymatic and non-enzymatic steps that lead to the generation of deaminated purine nucleotides are shown, as well as their removal by the inosine triphosphate pyrophosphatase (ITPA) and reintegration into canonical purine nucleotide metabolism or potential incorporation into nucleic acids. (d)ATP, (deoxy)adenosine triphosphate; (d)ITP, (deoxy)inosine triphosphate; (d)XTP, (deoxy)xanthosine triphosphate; (d)GTP, (deoxy)guanosine triphosphate; AMP, adenosine monophosphate; (d)IMP, (deoxy)inosine monophosphate; (d)XMP, (deoxy)xanthosine monophosphate; IMPDH, inosine monophosphate dehydrogenase; XMPP, xanthosine monophosphate phosphatase.

### Loss of ITPA leads to an upregulation of transcripts involved in biotic stress

We further investigated whether the loss of *ITPA* leading to the accumulation of (deoxy)inosine in nucleic acids and inosylates in nucleotide pools has an influence on the transcriptome. An RNAseq analysis was performed with RNA isolated from rosette leaves of 35-day-old wild type and *itpa-1* plants that had been grown side-by-side in a randomized fashion and for which we had already confirmed differential levels of inosine in RNA (Fig. 10b). Plants lacking ITPA showed an upregulation of 256 transcripts and a downregulation of only 28 transcripts at a log_2_ fold change (log_2_ FC) of ≥ 2 and ≤ −2 respectively in addition to a false discovery rate (FDR) of ≤ 0.05. Many regulated transcripts were involved in biological processes associated with stress responses and often encode proteins with kinase activity or transcription factors as reported by GO (gene ontology) enrichment analysis (Fig. S8). Upon closer inspection, we found that especially transcripts related to the salicylic acid response (Fig. 12a), systemic acquired resistance (SAR, Fig. 12b) and aging (Fig. 12c) were upregulated.

**Figure 12.**
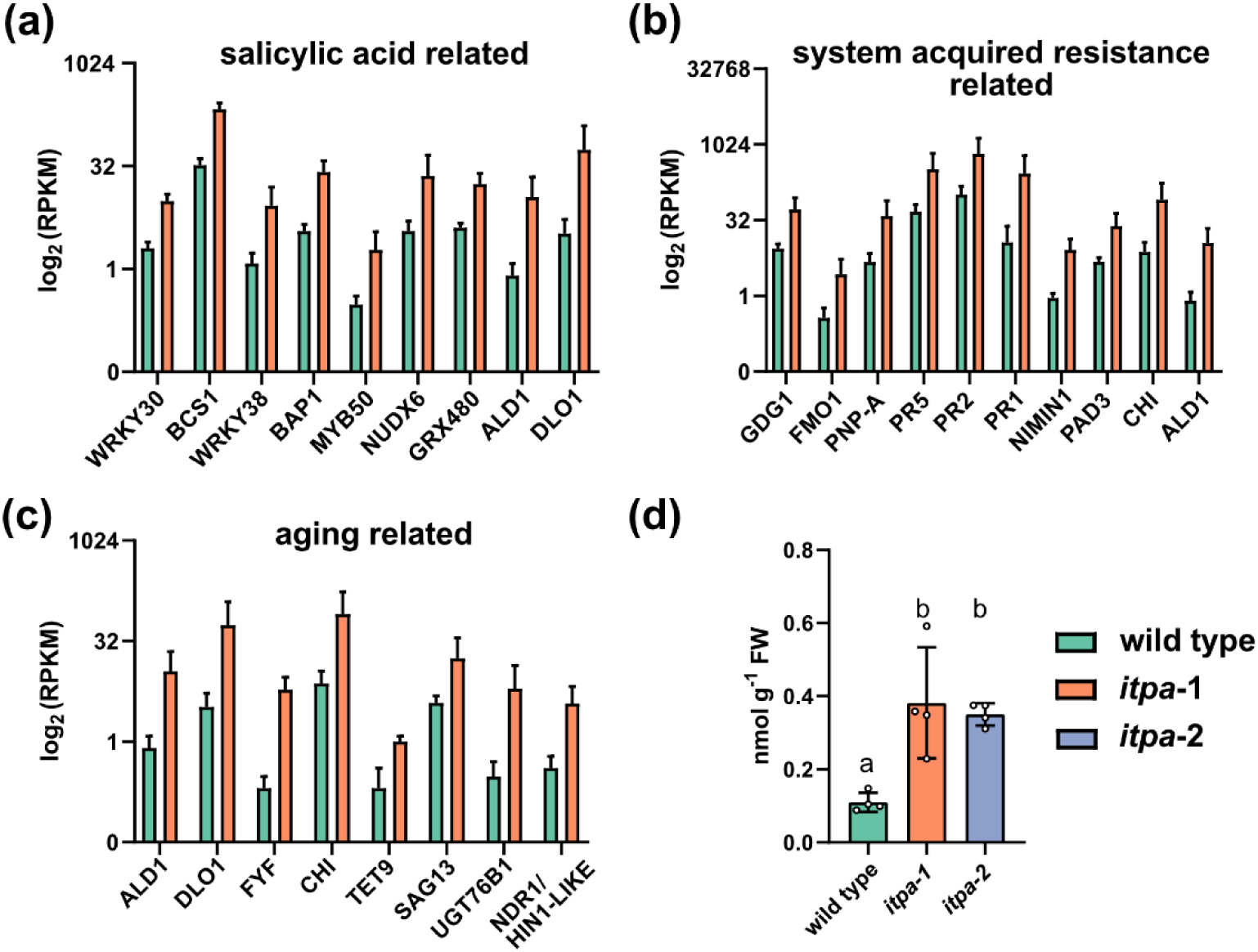
Loss-of-function of ITPA leads to differential upregulation of transcripts associated with salicylic acid, systemic acquired resistance as well as aging and increased endogenous concentrations of salicylic acid. (a) RNA was isolated from 35-day-old Arabidopsis rosettes and analyzed by RNAseq. Log_2_ (RPKM) values are shown for genes associated with response to salicylic acid (GO:0009751) and differentially regulated between wild type and *itpa-1* samples. Error bars are SD, *n* = 3. Every replicate represents a pool of two different plants. (b) As in (a) but for genes associated with systemic acquired resistance (GO:0009627). (c) As in (a) but for genes associated with aging (GO:0007568). (d) Concentrations of salicylic acid in 35-day-old Arabidopsis rosettes analyzed by LC-MS. Error bars are SD, *n* = 4. Every replicate represents a different plant. Statistical analysis was performed by a two-sided Tukey’s pairwise comparison using the sandwich variance estimator. Different letters indicate *p* values < 0.05. All *p* values can be found in the Table S4.

Most prominently transcripts of pathogenesis related 1 (At2g14610), pathogenesis related 2 (At3g57260), pathogenesis related 5 (At1g75040), NIM1 interacting 1 (At1g02450), senescence associated gene 13 (At2g29350), AGD2-like defense response protein 1 (At2g13810), DMR6-like oxygenase 1 (At4g10500) and UDP-dependent glycosyltransferase 76B1 (At3g11340) were differentially regulated. Since the transcriptional profiles suggested that the concentration of stress related phytohormones may be altered, we quantified salicylic acid (SA). Significantly higher SA concentrations were found in 35-day-old plants lacking ITPA compared to the corresponding wild type (Fig. 12d). The expression of *ITPA* itself seems not to be influenced by SA according to public transcriptomic data (Winter et al., 2007).

## DISCUSSION

Although the existence of ITPA in plants has been suggested based on sequence comparisons (Lin et al., 2001; James et al., 2021), these *in silico* predictions have never been experimentally tested nor has an *ITPA* mutant been characterized in any plant species. Here we demonstrate that plants have ITPA and that it is involved in the metabolite damage repair system in Arabidopsis. The kinetic properties of the enzyme from Arabidopsis are similar to those of the yeast ortholog HAM1p. Interestingly, the human ITPA has a 250-fold higher K_M_ and a 300 to 500-fold greater catalytic activity than the plant and yeast enzymes (Table 3) which might be an adaptation to high ITP concentrations in some human cells, for example erythrocytes (Behmanesh et al., 2009). However, in other human cells, the ITP concentration must be relatively low because it was generally not detectable (Sakumi et al., 2010).

The K_M_ of Arabidopsis ITPA for ITP of 7.1 µM is approximately 1000-fold higher than the cellular concentration of ITP (9.6 nM) determined in *itpa* background, which will be even lower in the wild type. We were unable to detect dITP but could show that mutation of *ITPA* led to more deoxyinosine in DNA. Thus, dITP is likely an *in vivo* substrate of ITPA (K_M_ = 1.7 µM for dITP). We estimate that the dITP concentration is in the low pM range assuming that the ratio of ITP to dITP is comparable to the ATP to dATP ratio of about 500 to 1000 (Straube et al., 2021). A difference of several orders of magnitude between K_M_ and cellular substrate concentration is also known for other house-keeping enzymes (Galperin et al., 2006; Sakumi et al., 2010; Heinemann et al., 2021).

XTP, dXTP and (deoxy)xanthosine in nucleic acids could not be detected, indicating that the deamination of (deoxy)guanosine in DNA and RNA and the phosphorylation of (d)XMP are very rare. Therefore, ITPA seems mainly involved with ITP and IDP and somewhat with dITP dephosphorylation *in vivo* (Fig. 11). The activity of Arabidopsis ITPA with IDP is interesting, since the yeast and human orthologs have a rather poor activity with IDP (Davies et al., 2012; Holmes et al., 1979). Human cells possess a nudix hydrolase (NUDT16) which hydrolyzes preferably IDP and dIDP, acting synergistically with ITPA to protect cells from high levels of (d)ITP (Abolhassani et al., 2010). This enzyme is missing in Arabidopsis, which may explain why the plant enzyme is also an efficient diphosphate hydrolase. IDP is likely the product of ADP deamination (Fig. 9) and ADP is also a substrate of the ribonucleotide reductase (RNR; Wang and Liu, 2006), an enzyme complex catalyzing the first committed step in the synthesis of deoxyribonucleotide triphosphates. Assuming that RNR accepts IDP as a substrate the reaction would lead to the formation of deoxyinosine diphosphate, which could be phosphorylated and then be incorporated into DNA.

According to our data, an important source of ITP is probably the deamination of ATP. The concentration of the adenylates is by far the highest of all nucleotides (Straube et al., 2021) and even a small rate of deamination will generate significant amounts of ITP. Additionally, this rate can be enhanced by oxidative stress, which can occur in plant cells as a result of abiotic and biotic stress. AMP kinases may also contribute to the IDP and ITP pool because they can use IMP as substrate albeit with lower activity than for AMP. The possibility of spurious phosphorylation by kinases involved in nucleotide metabolism has been discussed recently (Chen et al., 2018; Chen and Witte, 2020). However, *in vivo* there is far more AMP than IMP and our manipulation of IMP in *N. benthamiana* leaves showed no correlation of IMP and ITP concentrations (Fig. 7), indicating that the contribution of spurious IMP phosphorylation to the IDP and ITP pools is rather small.

It is surprising that ITPA is only located in the cytosol, nucleus and plastids but not in the mitochondria because ATP occurs in each of these compartments and mitochondria also perform nucleic acid biosynthesis. Additionally, the respiratory chain and photosystems are main sources for ROS in plant cells (Janků et al., 2019), potentially leading to nucleotide and nucleic acid damage, such as nucleotide deamination (Cadet and Wagner, 2013), in mitochondria and plastids, respectively (Tripathi et al., 2020). It is known that ATP and ADP are rapidly exchanged over the mitochondrial membrane because the mitochondria supply the cell with ATP produced by oxidative phosphorylation (Braun, 2020). Therefore, cytosolic ITPA may be able to also purify the mitochondrial adenylate pool – this however implies that ITP and IDP are also rapidly exchanged over the mitochondrial membrane (Haferkamp et al., 2011). By contrast, plastids exchange adenylates with the cytosol mainly at night for ATP import whereas during the day they internally generate ATP from ADP driven by photosynthesis for triosephosphate production. Plastids may require their own ITPA because in the light ROS production is high and adenylates are not exchanged with the cytosol.

Mutant plants lacking *ITPA* show a slightly earlier senescence and increased cell death (Fig. 4a-d). This is a relatively moderate phenotype compared to what occurs in mammals. Mice without functional ITPA die shortly after birth (Behmanesh et al., 2009). A null mutation of *ITPA* in humans leads to severe developmental abnormalities and is also lethal (Handley et al., 2019). Humans with missense mutations in *ITPA* develop severe side effects when treated with purine analoga like ribavirin or thiopurines (Simone et al., 2013). By contrast, the single cell eukaryote *S. cerevisiae* and also the model bacterium *E. coli* show no particular physiological phenotype when they lack ITPA (Clyman and Cunningham, 1987; Pavlov, 1986; Noskov et al., 1996b). Nonetheless, the *E. coli* mutant has an induced SOS response, a pathway that is turned on by DNA damage (Clyman and Cunningham, 1987).

So why do plants lacking ITPA activity show symptoms of early senescence? This may has to do with the accumulation of (deoxy)inosine in DNA and RNA in the mutant (Fig. 10, Fig. S6). Deaminated purines in RNA were shown to lead to reduced polymerase fidelity, RNA structure changes, altered stability and mistranslation (Thomas et al., 1998; Ji et al., 2017), and in case of DNA to mutations (Spee et al., 1993; DeVito et al., 2017) and genome instability (Yoshimura et al., 2007). We observed a higher amount of inosine in RNA of older *itpa* plants (Fig. 10 and Fig. 6) indicating that the problems may become more severe with age. DNA damage can lead to programmed cell death (PCD) in Arabidopsis (Wang and Liu, 2006), and we saw a higher incidence of PCD in the *itpa* background during senescence (Fig. 4d). An early senescence phenotype similar to that in *itpa* plants was also observed in plants with a higher frequency of unrepaired DNA double-strand breaks (Li et al., 2020). In summary, damage to DNA and RNA by aberrant nucleotide incorporation may be the cause for the earlier senescence and the increased PCD rate. Alternatively, the detrimental effects of *ITPA* mutation may also be unrelated to the alteration of nucleic acids but caused by ITP accumulation. ITP can directly influence cellular processes that either require or are inhibited by ATP or GTP (Vanderheiden, 1979; Burton et al., 2005; Osheroff et al., 1983; Klinker and Seifert, 1997). However, compared to the sizes of the ATP and GTP pools, the ITP pool is very small even in the *itpa* background, which argues against a direct metabolic interference of ITP.

The abundance of transcripts associated with salicylic acid (SA), senescence and systemic acquired resistance increases in *itpa* plants and they accumulate SA (Fig. 12 and Fig. S8). Mutants with constitutive high levels of SA are known to senesce prematurely. The higher SA levels in *itpa* plants may thus present an explanation for the early senescence phenotype observed (Fig. 4a-d) and may also affect plant-pathogen interactions. How the mutation of ITPA and the induction of SA are connected is unclear. We hypothesize that an accumulation of deoxyinosine in DNA causes more DNA repair and turnover associated with occasional double strand breaks. It is known that in plants with more unrepaired DNA double-strand breaks the concentration of SA is increased (Li et al., 2020). It also has been shown that SA can trigger DNA damage (Hadwiger and Tanaka, 2017) and that proteins involved in DNA damage response, like RAD51, interact synergistically to trigger increased transcription of genes involved in plant immunity (Yan et al., 2013). Taken together, it is possible that the transcripts involved in SA response, SAR and aging are upregulated as a consequence of DNA damage.

Another exciting possibility is that ITP or IDP are sensed directly and trigger a response. Such a signaling function has been shown for other nucleotides like extracellular ATP (Cao et al., 2014; Witte and Herde, 2020), NAD and NADH (Pétriacq et al., 2016; Fernie et al., 2020) or the recently discovered 2’,3’-cAMP (Yu et al., 2021) and is speculated for many other nucleotides like mononucleoside polyphosphates, dinucleoside polyphosphates or adenosine-phosphoramidate (Pietrowska-Borek et al., 2020).

Deamination of adenylates is an unavoidable chemical reaction and most organisms including plants have ITPA to protect their nucleic acids from the reaction products. Whether (d)ITP and (d)IDP are more than just an undesirable metabolic byproduct remains to be investigated.

## ACKNOWLEDGEMENT

We would like to express our gratitude to Holger Eubel and Björn Heinemann for providing the hydroponic system and André Specht for technical assistance. We also like to thank Anke Steppuhn (University of Hohenheim) for the donation of the phytohormone isotope standards. We furthermore like to express our gratitude to Sören Budig and Frank Schaarschmidt (Leibniz University, Hannover) for advice on the statistical analysis. We like to acknowledge the support by the Deutsche Forschungsgemeinschaft (HE 5949/4-1 to M.H.), (grant no. WI3411/7-1 and WI3411/8-1 to C-P.W.), and (grant no. INST 187/741-1 FUGG).

## AUTHOR CONTRIBUTIONS

HS and MH designed the research, HS, JS, JR and MN performed research, HS, JS, JR and MH analyzed the data and HS, CPW and MH interpreted the data. HS, CPW and MH wrote the manuscript, all authors read and revised the manuscript and agreed on the final version.

## DATA AVAILABILLITY

The data that supports the findings of this study are available in the supporting information of this article

Information about the mutant lines in this article can be obtained in the GenBank/EMBL data libraries under the following accession number:

At4g13720 (ITPA).

Raw read data from Illumina sequencing is deposited at the Sequence Read Archive (SRA) on NCBI under following accession number: PRJNA807491.

## SUPPORTING INFORMATION

**Table S1.** List of primers used in this study

**Table S2.** List of differentially expressed genes

**Table S3.** List of the GO term enrichment analysis

**Table S4.** Raw and statistic data

**Table S5.** Substrates tested for biochemical activity with ITPA

**Table S6.** Ratios (deoxy)inosine per million molecules (deoxy)adenosine in DNA and RNA, respectively

**Figure S1.** Comparative sequence analysis of inosine triphosphate pyrophosphatases from several phylogenetically distant model organisms of all kingdoms of life

**Figure S2.** Comparative sequence analysis of inosine triphosphate pyrophosphatases

**Figure S3.** Phenotyping of wild type and *itpa* mutant plants

**Figure S4.** Chromatograms of ITP and IDP analyzed by hypercarb chromatography

**Figure S5.** Relative concentration of different nucleotides in rosette leaves of 35-day-old wild type and itpa mutant plants

**Figure S6.** Manipulation of IMP content in leaves of *N. benthamiana* alters the concentration of GMP and XMP, and in tendency also GTP

**Figure S7.** Concentration of inosine in isolated RNA of wild type and itpa-1 plants

**Figure S8.** Loss-of-function of ITPA leads to differential upregulation of transcripts associated with GO:terms of biological processes, molecular functions and cellular components

**Supporting methods**

## REFERENCES

Abolhassani, N., Iyama, T., Tsuchimoto, D., Sakumi, K., Ohno, M., Behmanesh, M., and Nakabeppu, Y. (2010). NUDT16 and ITPA play a dual protective role in maintaining chromosome stability and cell growth by eliminating dIDP/IDP and dITP/ITP from nucleotide pools in mammals. Nucleic Acids Research 38 (9): 2891– 2903.

Alonso, J.M., Stepanova, A.N., Leisse, T.J., Kim, C.J., Chen, H., Shinn, P., Stevenson, D.K., Zimmerman, J., Barajas, P., Cheuk, R., Gadrinab, C., Heller, C., Jeske, A., Koesema, E., Meyers, C.C., Parker, H., Prednis, L., Ansari, Y., Choy, N., Deen, H., Geralt, M., Hazari, N., Hom, E., Karnes, M., Mulholland, C., Ndubaku, R., Schmidt, I., Guzman, P., Aguilar-Henonin, L., Schmid, M., Weigel, D., Carter, D.E., Marchand, T., Risseeuw, E., Brogden, D., Zeko, A., Crosby, W.L., Berry, C.C., and Ecker, J.R. (2003). Genome-wide insertional mutagenesis of Arabidopsis thaliana. Science 301 (5633): 653–657.

Andrews, S. (2010). FastQC: A Quality Control Tool for High Throughput Sequence Data. Available online at: http://www.bioinformatics.babraham.ac.uk/projects/fastqc/

Baccolini, C., and Witte, C.-P. (2019). AMP and GMP catabolism in arabidopsis converge on xanthosine, which is degraded by a nucleoside hydrolase heterocomplex. The Plant Cell 31 (3): 734–751.

Batista-Silva, W., Heinemann, B., Rugen, N., Nunes-Nesi, A., Araújo, W.L., Braun, H.-P., and Hildebrandt, T.M. (2019). The role of amino acid metabolism during abiotic stress release. Plant, Cell & Environment 42 (5): 1630–1644.

Behmanesh, M., Sakumi, K., Abolhassani, N., Toyokuni, S., Oka, S., Ohnishi, Y.N., Tsuchimoto, D., and Nakabeppu, Y. (2009). ITPase-deficient mice show growth retardation and die before weaning. Cell death and differentiation 16 (10): 1315–1322.

Bolte, S. and Cordelieres, F.P. (2006). A guided tour into subcellular colocalization analysis in light microscopy. Journal of Microscopy, 224 (3): 213–232

Braun, H.-P. (2020). The oxidative phosphorylation system of the mitochondria in plants. Mitochondrion 53: 66–75.

Budke, B., and Kuzminov, A. (2006). Hypoxanthine incorporation is nonmutagenic in Escherichia coli. Journal of Bacteriology 188 (18): 6553–6560.

Budke, B., and Kuzminov, A. (2010). Production of clastogenic DNA precursors by the nucleotide metabolism in Escherichia coli. Molecular Microbiology 75 (1): 230–245.

Burgis, N.E., Brucker, J.J., and Cunningham, R.P. (2003). Repair system for noncanonical purines in Escherichia coli. Journal of Bacteriology 185 (10): 3101–3110.

Burton, K., White, H., and Sleep, J. (2005). Kinetics of muscle contraction and actomyosin NTP hydrolysis from rabbit using a series of metal-nucleotide substrates. The Journal of Physiology 563 (Pt 3): 689–711.

Cadet, J., and Wagner, J.R. (2013). DNA base damage by reactive oxygen species, oxidizing agents, and UV radiation. Perspectives in Biology 5 (2).

Cao, Y., Tanaka, K., Nguyen, C.T., and Stacey, G. (2014). Extracellular ATP is a central signaling molecule in plant stress responses. Current Opinion in Plant Biology 20: 82– 87.

Carnelli, A., Michelis, M.I. de, and Rasi-Caldogno, F. (1992). Plasma Membrane Ca-ATPase of Radish Seedlings I. Biochemical characterization using ITP as a substrate. Plant Physiology 98 (3): 1196–1201.

Carole L Linster, Emile Van Schaftingen, and Andrew D Hanson (2013). Metabolite damage and its repair or pre-emption. Nature Chemical Biology 9 (2): 72–80.

Chen, M., Urs, M.J., Sánchez-González, I., Olayioye, M.A., Herde, M., and Witte, C.-P. (2018). m6A RNA degradation products are catabolized by an evolutionarily conserved N6-Methyl-AMP deaminase in plant and mammalian cells. The Plant Cell 30 (7): 1511–1522.

Chen, M., and Witte, C.-P. (2020). A kinase and a glycosylase catabolize pseudouridine in the peroxisome to prevent toxic pseudouridine monophosphate accumulation. The Plant Cell 32 (3): 722–739.

Cho, U.-H., and Seo, N.-H. (2005). Oxidative stress in Arabidopsis thaliana exposed to cadmium is due to hydrogen peroxide accumulation. Plant Science 168 (1): 113–120.

Claus-Peter Witte, Laurent Noël, Janine Gielbert, Jane Parker, and Tina Romeis (2004). Rapid one-step protein purification from plant material using the eight-amino acid StrepII epitope. Plant Molecular Biology 55 (1): 135–147.

Clyman, J., and Cunningham, R.P. (1987). Escherichia coli K-12 mutants in which viability is dependent on recA function. Journal of Bacteriology 169 (9): 4203–4210.

Corpas, F.J., Leterrier, M., Valderrama, R., Airaki, M., Chaki, M., Palma, J.M., and Barroso, J.B. (2011). Nitric oxide imbalance provokes a nitrosative response in plants under abiotic stress. Plant Science 181 (5): 604–611.

Crécy-Lagard, V. de, Haas, D., and Hanson, A.D. (2018). Newly-discovered enzymes that function in metabolite damage-control. Current Opinion in Chemical Biology 47: 101–108.

Dahncke, K., and Witte, C.-P. (2013). Plant purine nucleoside catabolism employs a guanosine deaminase required for the generation of xanthosine in Arabidopsis. The Plant Cell 25 (10): 4101–4109.

Davies, O., Mendes, P., Smallbone, K., and Malys, N. (2012). Characterisation of multiple substrate-specific (d)ITP/(d)XTPase and modelling of deaminated purine nucleotide metabolism. BMB Reports 45 (4): 259–264.

Demidchik, V. (2015). Mechanisms of oxidative stress in plants: From classical chemistry to cell biology. Environmental and Experimental Botany 109: 212–228.

DeVito, S., Woodrick, J., Song, L., and Roy, R. (2017). Mutagenic potential of hypoxanthine in live human cells. Mutation Research 803–805: 9–16.

Di Sanità Toppi, L., and Gabbrielli, R. (1999). Response to cadmium in higher plants. Environmental and Experimental Botany 41 (2): 105–130.

Dobrzanska, M., Szurmak, B., Wyslouch-Cieszynska, A., and Kraszewska, E. (2002). Cloning and characterization of the first member of the Nudix family from Arabidopsis thaliana. Journal of Biological Chemistry 277 (52): 50482–50486.

Dubois, E., Córdoba-Cañero, D., Massot, S., Siaud, N., Gakière, B., Domenichini, S., Guérard, F., Roldan-Arjona, T., and Doutriaux, M.-P. (2011). Homologous recombination is stimulated by a decrease in dUTPase in Arabidopsis. PLoS ONE 6 (4): e18658.

Fernie, A., Hashida, S.-N., Yoshimura, K., Gakière, B., Mou, Z., and Pétriacq, P. (2020). Editorial: NAD Metabolism and Signaling in Plants. Frontiers in Plant Science 11: 146.

Fraser, J.H., Meyers, H., Henderson, J.F., Brox, L.W., and McCoy, E.E. (1975). Individual variation in inosine triphosphate accumulation in human erythrocytes. Clinical Biochemistry 8 (1-6): 353–364.

Frederico, L.A., Kunkel, T.A., and Shaw, B.R. (1990). A sensitive genetic assay for the detection of cytosine deamination: determination of rate constants and the activation energy. Biochemistry 29 (10): 2532–2537.

Gall, A.D., Gall, A., Moore, A.C., Aune, M.K., Heid, S., Mori, A., and Burgis, N.E. (2013). Analysis of human ITPase nucleobase specificity by site-directed mutagenesis. Biochimie 95 (9): 1711–1721.

Hadwiger, L.A., and Tanaka, K. (2017). Non-host resistance: DNA damage is associated with sa signaling for induction of pr genes and contributes to the growth suppression of a pea pathogen on pea endocarp tissue. Frontiers in Plant Science 8: 446.

Haferkamp, I., Fernie, A.R., and Neuhaus, H.E. (2011). Adenine nucleotide transport in plants: much more than a mitochondrial issue. Trends in Plant Science 16 (9): 507– 515.

Handley, M.T., Reddy, K., Wills, J., Rosser, E., Kamath, A., Halachev, M., Falkous, G., Williams, D., Cox, P., Meynert, A., Raymond, E.S., Morrison, H., Brown, S., Allan, E., Aligianis, I., Jackson, A.P., Ramsahoye, B.H., Kriegsheim, A. von, Taylor, R.W., Finch, A.J., and FitzPatrick, D.R. (2019). ITPase deficiency causes a Martsolf-like syndrome with a lethal infantile dilated cardiomyopathy. PLoS Genetics 15 (3): e1007605.

Hanson, A.D., Henry, C.S., Fiehn, O., and Crécy-Lagard, V. de (2016). Metabolite damage and metabolite damage control in plants. Annual Review of Plant Biology 67: 131–152.

Harris, C., and Baulcombe, D. (2015). Chlorophyll content assay to quantify the level of necrosis induced by different R gene/elicitor combinations after transient expression. Bio-Protocol 5 (23).

Heinemann, K.J., Yang, S.-Y., Straube, H., Medina-Escobar, N., Varbanova-Herde, M., Herde, M., Rhee, S., and Witte, C.-P. (2021). Initiation of cytosolic plant purine nucleotide catabolism involves a monospecific xanthosine monophosphate phosphatase. Nature communications 12 (1): 6846.

Holmes, S.L., Turner, B.M., and Hirschhorn, K. (1979). Human inosine triphosphatase: Catalytic properties and population studies. Clinica Chimica Acta 97 (2-3): 143–153.

Hooper, C.M., Castleden, I.R., Tanz, S.K., Aryamanesh, N., and Millar, A.H. (2017). SUBA4: the interactive data analysis centre for Arabidopsis subcellular protein locations. Nucleic Acids Research 45 (D1): D1064–D1074.

James, A.M., Seal, S.E., Bailey, A.M., and Foster, G.D. (2021). Viral inosine triphosphatase: A mysterious enzyme with typical activity, but an atypical function. Molecular Plant Pathology 22 (3): 382–389.

Janků, M., Luhová, L., and Petřivalský, M. (2019). On the origin and fate of reactive oxygen species in plant cell compartments. Antioxidants (Basel, Switzerland) 8 (4).

Ji, D., Stepchenkova, E.I., Cui, J., Menezes, M.R., Pavlov, Y.I., and Kool, E.T. (2017). Measuring deaminated nucleotide surveillance enzyme ITPA activity with an ATP-releasing nucleotide chimera. Nucleic Acids Research 45 (20): 11515–11524.

Jiang, H.-P., Xiong, J., Liu, F.-L., Ma, C.-J., Tang, X.-L., Yuan, B.-F., and Feng, Y.-Q. (2018). Modified nucleoside triphosphates exist in mammals. Chemical Science 9 (17): 4160–4167.

Kamiya, H. (2003). Mutagenic potentials of damaged nucleic acids produced by reactive oxygen/nitrogen species: approaches using synthetic oligonucleotides and nucleotides: survey and summary. Nucleic Acids Research 31 (2): 517–531.

Keya, C.A., Crozier, A., and Ashihara, H. (2003). Inhibition of caffeine biosynthesis in tea (Camellia sinensis) and coffee (Coffea arabica) plants by ribavirin. FEBS Letters 554 (3): 473–477.

Klinker, J.F., and Seifert, R. (1997). Functionally nonequivalent interactions of guanosine 5′-triphosphate, inosine 5′-triphosphate, and xanthosine 5′-triphosphate with the retinal G-protein, transducin, and with Gi-proteins in HL-60 leukemia cell membranes. Biochemical Pharmacology 54 (5): 551–562.

Kong, R.P.W., Siu, S.O., Lee, S.S.M., Lo, C., and Chu, I.K. (2011). Development of online high-/low-pH reversed-phase-reversed-phase two-dimensional liquid chromatography for shotgun proteomics: a reversed-phase-strong cation exchange-reversed-phase approach. Journal of Chromatography. A 1218 (23): 3681–3688.

Kuskovsky, R., Buj, R., Xu, P., Hofbauer, S., Doan, M.T., Jiang, H., Bostwick, A., Mesaros, C., Aird, K.M., and Snyder, N.W. (2019). Simultaneous isotope dilution quantification and metabolic tracing of deoxyribonucleotides by liquid chromatography high resolution mass spectrometry. Analytical Biochemistry 568: 65–72.

Leiva, A.M., Jimenez, J., Sandoval, H., Perez, S., and Cuellar, W.J. (2022). Complete genome sequence of a novel secovirid infecting cassava in the Americas. Archives of Virology.

Lerma-Ortiz, C., Jeffryes, J.G., Cooper, A.J.L., Niehaus, T.D., Thamm, A.M.K., Frelin, O., Aunins, T., Fiehn, O., Crécy-Lagard, V. de, Henry, C.S., and Hanson, A.D. (2016). ‘Nothing of chemistry disappears in biology’: the Top 30 damage-prone endogenous metabolites. Biochemical Society Transactions 44 (3): 961–971.

Li, Z., Kim, J.H., Kim, J., Lyu, J.I., Zhang, Y., Guo, H., Nam, H.G., and Woo, H.R. (2020). ATM suppresses leaf senescence triggered by DNA double-strand break through epigenetic control of senescence-associated genes in Arabidopsis. The New Phytologist 227 (2): 473–484.

Lin, S., McLennan, A.G., Ying, K., Wang, Z., Gu, S., Jin, H., Wu, C., Liu, W., Yuan, Y., Tang, R., Xie, Y., and Mao, Y. (2001). Cloning, expression, and characterization of a human inosine triphosphate pyrophosphatase encoded by the itpa gene. Journal of Biological Chemistry 276 (22): 18695–18701.

Lin, A.-J., Zhang, X.-H., Chen, M.-M., and Cao, Q. (2007). Oxidative stress and DNA damages induced by cadmium accumulation. Journal of Environmental Sciences 19 (5): 596–602.

Lindahl, T. (1993). Instability and decay of the primary structure of DNA. Nature 362 (6422): 709–715.

Lindahl, T., and Nyberg, B. (1974). Heat-induced deamination of cytosine residues in deoxyribonucleic acid. Biochemistry 13 (16): 3405–3410.

Littlejohn, G.R., Gouveia, J.D., Edner, C., Smirnoff, N., and Love, J. (2010). Perfluorodecalin enhances in vivo confocal microscopy resolution of Arabidopsis thaliana mesophyll. The New Phytologist 186 (4): 1018–1025.

Liu, S., Yu, F., Yang, Z., Wang, T., Xiong, H., Chang, C., Yu, W., and Li, N. (2018). Establishment of Dimethyl Labeling-based Quantitative Acetylproteomics in Arabidopsis. Molecular & Cellular Proteomics 17 (5): 1010–1027.

Galperin, M. Y., Moroz, O. V., Wilson, K. S., and Murzin, A. G. (2006). House cleaning, a part of good housekeeping. Molecular Microbiology 59 (1): 5–19.

Muraoka, M., Fukuzawa, H., Nishida, A., Okano, K., Tsuchihara, T., Shimoda, A., Suzuki, Y., Sato, M., Osumi, M., and Sakai, H. (1999). The effects of various GTP analogues on microtubule assembly. Cell Structure and Function 24 (2): 101–109.

Myrach, T., Zhu, A., and Witte, C.-P. (2017). The assembly of the plant urease activation complex and the essential role of the urease accessory protein G (UreG) in delivery of nickel to urease. The Journal of Biological Chemistry 292 (35): 14556–14565.

Nagy, G.N., Leveles, I., and Vértessy, B.G. (2014). Preventive DNA repair by sanitizing the cellular (deoxy)nucleoside triphosphate pool. The FEBS journal 281 (18): 4207– 4223.

Niehaus, M., Straube, H., Künzler, P., Rugen, N., Hegermann, J., Giavalisco, P., Eubel, H., Witte, C.-P., and Herde, M. (2020). Rapid Affinity Purification of Tagged Plant Mitochondria (Mito-AP) for Metabolome and Proteome Analyses. Plant Physiology 182 (3): 1194–1210.

Niehaus, T.D., Patterson, J.A., Alexander, D.C., Folz, J.S., Pyc, M., MacTavish, B.S., Bruner, S.D., Mullen, R.T., Fiehn, O., and Hanson, A.D. (2019). The metabolite repair enzyme Nit1 is a dual-targeted amidase that disposes of damaged glutathione in Arabidopsis. The Biochemical Journal 476 (4): 683–697.

Niehaus, T.D., Richardson, L.G.L., Gidda, S.K., ElBadawi-Sidhu, M., Meissen, J.K., Mullen, R.T., Fiehn, O., and Hanson, A.D. (2014). Plants utilize a highly conserved system for repair of NADH and NADPH hydrates. Plant Physiology 165 (1): 52–61.

Noskov, V.N., Staak, K., Shcherbakova, P.V., Kozmin, S.G., Negishi, K., Ono, B.-C., Hayatsu, H., and Pavlov, Y.I. (1996). HAM1, the gene controlling 6-N-hydroxylaminopurine sensitivity and mutagenesis in the yeast Saccharomyces cerevisiae. Yeast 12 (1): 17–29.

Fernández-Bautista, N., Domínguez-Núñez, J. A., Moreno, M. M. C., and Berrocal-Lobo, M. (2016). Plant Tissue Trypan Blue Staining During Phytopathogen Infection. Bio-Protocol 6 (24): e2078–e2078.

Oberacker, P., Stepper, P., Bond, D.M., Höhn, S., Focken, J., Meyer, V., Schelle, L., Sugrue, V.J., Jeunen, G.-J., Moser, T., Hore, S.R., Meyenn, F. von, Hipp, K., Hore, T.A., and Jurkowski, T.P. (2019). Bio-On-Magnetic-Beads (BOMB): Open platform for high-throughput nucleic acid extraction and manipulation. PLoS Biology 17 (1): e3000107.

Osheroff, N., Shelton, E.R., and Brutlag, D.L. (1983). DNA topoisomerase II from Drosophila melanogaster. Relaxation of supercoiled DNA. Journal of Biological Chemistry 258 (15): 9536–9543.

Pang, B., McFaline, J.L., Burgis, N.E., Dong, M., Taghizadeh, K., Sullivan, M.R., Elmquist, C.E., Cunningham, R.P., and Dedon, P.C. (2012). Defects in purine nucleotide metabolism lead to substantial incorporation of xanthine and hypoxanthine into DNA and RNA. Proceedings of the National Academy of Sciences of the United States of America 109 (7): 2319–2324.

Pétriacq, P., Ton, J., Patrit, O., Tcherkez, G., and Gakière, B. (2016). NAD Acts as an Integral Regulator of Multiple Defense Layers. Plant Physiology 172 (3): 1465–1479.

Pietrowska-Borek, M., Dobrogojski, J., Sobieszczuk-Nowicka, E., and Borek, S. (2020). New Insight into Plant Signaling: Extracellular ATP and Uncommon Nucleotides. Cells 9 (2).

Rampazzo, C., Miazzi, C., Franzolin, E., Pontarin, G., Ferraro, P., Frangini, M., Reichard, P., and Bianchi, V. (2010). Regulation by degradation, a cellular defense against deoxyribonucleotide pool imbalances. Mutation Research 703 (1): 2–10.

Rinne, J., Witte, C.-P., and Herde, M. (2021). Loss of MAR1 Function is a Marker for Co-Selection of CRISPR-Induced Mutations in Plants. Frontiers in Genome Editing 3: 723384.

Rudd, S.G., Valerie, N.C.K., and Helleday, T. (2016). Pathways controlling dNTP pools to maintain genome stability. DNA Repair 44: 193–204.

Sakumi, K., Abolhassani, N., Behmanesh, M., Iyama, T., Tsuchimoto, D., and Nakabeppu, Y. (2010). ITPA protein, an enzyme that eliminates deaminated purine nucleoside triphosphates in cells. Mutation Research 703 (1): 43–50.

Savchenko, A., Proudfoot, M., Skarina, T., Singer, A., Litvinova, O., Sanishvili, R., Brown, G., Chirgadze, N., and Yakunin, A.F. (2007). Molecular basis of the antimutagenic activity of the house-cleaning inosine triphosphate pyrophosphatase RdgB from Escherichia coli. Journal of Molecular Biology 374 (4): 1091–1103.

Simone, P.D., Pavlov, Y.I., and Borgstahl, G.E.O. (2013). ITPA (inosine triphosphate pyrophosphatase): from surveillance of nucleotide pools to human disease and pharmacogenetics. Mutation Research 753 (2): 131–146.

Spee, J.H., Vos, W.M. de, and Kuipers, O.P. (1993). Efficient random mutagenesis method with adjustable mutation frequency by use of PCR and dITP. Nucleic Acids Research 21 (3): 777–778.

Stenmark, P., Kursula, P., Flodin, S., Gräslund, S., Landry, R., Nordlund, P., and Schüler, H. (2007). Crystal structure of human inosine triphosphatase. Substrate binding and implication of the inosine triphosphatase deficiency mutation P32T. Journal of Biological Chemistry 282 (5): 3182–3187.

Straube, H., Niehaus, M., Zwittian, S., Witte, C.-P., and Herde, M. (2021). Enhanced nucleotide analysis enables the quantification of deoxynucleotides in plants and algae revealing connections between nucleoside and deoxynucleoside metabolism. The Plant Cell 33 (2): 270–289

Thomas, M.J., Platas, A.A., and Hawley, D.K. (1998). Transcriptional Fidelity and Proofreading by RNA Polymerase II. Cell 93 (4): 627–637.

Tian, T., Liu, Y., Yan, H., You, Q., Yi, X., Du, Z., Xu, W., and Su, Z. (2017). agriGO v2.0: a GO analysis toolkit for the agricultural community, 2017 update. Nucleic Acids Research 45 (W1): W122–W129.

Tomlinson, K.R., Pablo-Rodriguez, J.L., Bunawan, H., Nanyiti, S., Green, P., Miller, J., Alicai, T., Seal, S.E., Bailey, A.M., and Foster, G.D. (2019). Cassava brown streak virus Ham1 protein hydrolyses mutagenic nucleotides and is a necrosis determinant. Molecular Plant Pathology 20 (8): 1080–1092.

Traube, F. R., Schiffers, S., Iwan, K., Kellner, S., Spada, F., Müller, M., and Carell, T. (2019). Isotope-dilution mass spectrometry for exact quantification of noncanonical DNA nucleosides. Nature Protocols 14 (1): 283–312.

Tripathi, D., Nam, A., Oldenburg, D.J., and Bendich, A.J. (2020). Reactive Oxygen Species, Antioxidant Agents, and DNA Damage in Developing Maize Mitochondria and Plastids. Frontiers in Plant Science 11: 596.

Valderrama, R., Corpas, F.J., Carreras, A., Fernández-Ocaña, A., Chaki, M., Luque, F., Gómez-Rodríguez, M.V., Colmenero-Varea, P., Del Río, L.A., and Barroso, J.B. (2007). Nitrosative stress in plants. FEBS Letters 581 (3): 453–461.

Vanderheiden, B.S. (1969). Genetic studies of human erythrocyte inosine triphosphatase. Biochemical Genetics 3 (3): 289–297.

Vanderheiden, B.S. (1979). Possible implication of an inosinetriphosphate metabolic error and glutamic acid decarboxylase in paranoid schizophrenia. Biochemical Medicine 21 (1): 22–32.

Wang, C., and Liu, Z. (2006). Arabidopsis ribonucleotide reductases are critical for cell cycle progression, DNA damage repair, and plant development. The Plant Cell 18 (2): 350–365.

Weber, J., and Senior, A.E. (2001). Bi-site catalysis in F1-ATPase: does it exist? Journal of Biological Chemistry 276 (38): 35422–35428.

Winter, D., Vinegar, B., Nahal, H., Ammar, R., Wilson, G. V., Provart, N. J. (2007). An “Electronic Fluorescent Pictograph” Browser for Exploring and Analyzing Large-Scale Biological Data Sets. PLoS One 2(8): e718.

Witte, C.-P., and Herde, M. (2020). Nucleotide Metabolism in Plants. Plant Physiology 182 (1): 63–78.

Witte, C.-P., Rosso, M.G., and Romeis, T. (2005). Identification of three urease accessory proteins that are required for urease activation in Arabidopsis. Plant Physiology 139 (3): 1155–1162.

Yan, S., Wang, W., Marqués, J., Mohan, R., Saleh, A., Durrant, W.E., Song, J., and Dong, X. (2013). Salicylic acid activates DNA damage responses to potentiate plant immunity. Molecular Cell 52 (4): 602–610.

Yang, Z.-B., He, C., Ma, Y., Herde, M., and Ding, Z. (2017). Jasmonic Acid Enhances Al-Induced Root Growth Inhibition. Plant Physiology 173 (2): 1420–1433.

Yoneshima, Y., Abolhassani, N., Iyama, T., Sakumi, K., Shiomi, N., Mori, M., Shiomi, T., Noda, T., Tsuchimoto, D., and Nakabeppu, Y. (2016). Deoxyinosine triphosphate induces MLH1/PMS2- and p53-dependent cell growth arrest and DNA instability in mammalian cells. Scientific Reports 6: 32849.

Yoshimura, K., Ogawa, T., Ueda, Y., and Shigeoka, S. (2007). AtNUDX1, an 8-oxo-7,8-dihydro-2’-deoxyguanosine 5’-triphosphate pyrophosphohydrolase, is responsible for eliminating oxidized nucleotides in Arabidopsis. Plant & Cell Physiology 48 (10): 1438– 1449.

Yu, D., Song, W., Tan, E.Y.J., Liu, L., Cao, Y., Jirschitzka, J., Li, E., Logemann, E., Xu, C., Huang, S., Jia, A., Chang, X., Han, Z., Wu, B., Schulze-Lefert, P., and Chai, J. (2021). TIR domains of plant immune receptors are 2′,3′-cAMP/cGMP synthetases mediating cell death. bioRxiv

