## Supplementary material for "An inosine triphosphate pyrophosphatase safeguards plant nucleic acids from aberrant purine nucleotides": Figure S

```
Gly max   1 ---------------------------MAAARGLVLSRPVTFVTANAKKLEEVRAIL--G
Pha vul   1 MFINIDEYGLILRRSALLQKNQTEIWEMAAARGLLLSRPVTFVTGNAKKLEEVRAIL--G
Med tru   1 -----------------------MAAIKSAASGLVLPRPVTFVTGNAKKLEEVRAIL--G
Ara tha   1 ------------------------MAAAAAKAAVVLPRPVTFVTGNAKKLEEVKAII--G
Sol tub   1 ----------------------MAAAARTVGSGVVLPRSVTFVTGNAKKLEEVRAIL--G
Sol lyc   1 ---------------------MAAAAARTVGSGVVLPRSVTFVTGNAKKLEEVRTIL--G
Tri aes   1 ------------------------MSGAAAAVARVLPKAVTFVTGNAKKLEEVRAIL--G
Zea may   1 ----------------------------MSAAARVLPKAVTFVTGNAKKLEEVRAIL--G
Ory sat   1 ---------------------------MSGAAARALPKAVTFVTGNAKKLEEVRAIL--G
Mar pol   1 --------------------------MSAMGVRMQLSSPVTFVTGNANKLEEVKMIL--G
Sel moe   1 -------------------------AMALVRAEVVLKKPVTFVTGNAKKLEEVKMIL--G
Phy pat   1 --------------------------MAAMKVQMTLTKPVTFVTGNAKKLEEVKMIL--G
Dro mel   1 -----------------------------------MSKPITFVTGNAKKLEELVAIL--G
Hom sap   1 ------------------------------MAASLVGKKIVFVTGNAKKLEEVVQIL--G
Sus scr   1 ------------------------------MANSLVGKKIVFVTGNAKKLEEVIQIL--G
Xen lae   1 -------------------------------MAAAAGRSIVFVTGNAKKLEEVVQIL--G
Dan rer   1 -------------------------------MAVPTGRALVFVTGNAKKLEEVVQIL--G
Gal gal   1 -------------------------------MAAPVRRSVVFVTGNAKKLEEVTQIL--G
Cae ele   1 ----------------------------------MSLRKINFVTGNVKKLEEVKAIL---
Vol car   1 ---------------------------------MATPKKIYFATGNKKKLEEVTAILQSG
Chl rei   1 ---------------------------------MTTPSKIHFATGNKKKLEEVNAILAAG
Bot cin   1 ---------------------------------MAPPKTLNFITGNKNKLIEVKAIL--G
Neu cra   1 ----------------------------XMSAPSQARHIVNFITGNANKLGEVKAIL--E
Asp nig   1 ---------------------------------MASLEKLNFITGNKNKLAEVKAIL--G
Ust may   1 ---------------------------------XMSNPTLTFVTGNANKLREVQQIFSLT
Sac cer   1 ----------------------------------MSNNEIVFVTGNANKLKEVQSIL--T
Met jan   1 -------------------------------------MKIYFATGNPNKIKEANIIL--K
Bac sub   1 -----------------------------------XMKEAIIATHNPGKVKEFKEIL---
Dei rad   1 ------------------------XMQQQTGGRRRQIRRVVVATSNAGKVRELQGAL---
Esc col   1 ------------------------------------MQKVVLATGNAGKVRELASLL---

Gly max  32 NSIPF--QSLK------LDLPELQG-EPEDISKEKARIAA--LQVNGPVLVEDTCLCFNA
Pha vul  59 NSIPF--QSLK------LDLPELQG-EPEDISKEKARMAA--LQVNGPVLVEDTCLCFNA
Med tru  36 HSIPF--QSLK------LDLPELQG-EPEDISKEKARLAA--IQVKGPVLVEDTCLCFNA
Ara tha  35 NSIPF--KSLK------LDLPELQG-EPEDISKEKARLAA--LQVDGPVLVEDTCLCFNA
Sol tub  37 QSIPF--QSLK------LDLPELQG-EPEDISKEKARIAA--KEVNGPVLVEDTCLCFNA
Sol lyc  38 QSIPF--QSLK------LDLPELQG-EPEDISKEKARIAA--KEVNGPVLVEDTCLCFNA
Tri aes  35 SSIPF--QSLK------LDLPELQG-EPEDISKEKARMAA--SQVNGPVLVEDTCLCFNA
Zea may  31 SSVPF--QSLK------LDLPELQG-EPEYISKEKARIAA--SQVNGPVLVEDTCLCFNA
Ory sat  32 SSIPF--QSLK------LDLPELQG-EPEDISKEKARMAA--SQVNGPVLVEDTCLCFNA
Mar pol  33 QTIPF--RSVK------LDLPELQG-EPEDISKEKCRLAA--KEVNGPVLVEDTCLCFNA
Sel moe  34 NSIPF--STLR------VDLPELQG-EPEEISKEKARIAA--KQIDGAVLVEDTCLCFNA
Phy pat  33 QSIPF--QSLK------LDLPELQG-EPEDISKEKARLAA--KEIGGPVLVEDTCLCFNA
Dro mel  24 PSFPRTIVSKK------IDLPELQG-DIDEIAIKKCKEAA--RQVNGPVLVEDTSLCFNA
Hom sap  29 DKFPCTLVAQK------IDLPEYQG-EPDEISIQKCQEAV--RQVQGPVLVEDTCLCFNA
Sus scr  29 DKFPCTLVAQK------IDLPEYQG-EPDEISIQKCKEAA--RQVQGPVLVEDTCLCFNA
Xen lae  28 DKFPCKLVAKK------IDLPEYQG-EPDEISIQKCREAA--KQIQGPVIVEDTCLCFNA
Dan rer  28 DKFPYKLISKK------IDLPEYQG-EPDDISIQKCKEAA--RQVDGPVLVEDTCLCFRA
Gal gal  28 DSSPYTLVARK------IDLPEYQG-EPDEISVQKCREAA--RQIRGPVIVEDTCLCFNA
Cae ele  24 --KNFEVSNVD------VDLDEFQG-EPEFIAERKCREAV--EAVKGPVLVEDTSLCFNA
Vol car  28 APLPFVMEAVK------LDLPELQG-EPEEISKEKCRIAA--KLVGGAVMVEDTSLCFNA
Chl rei  28 AELPFEVVAAK------LDLPELQG-EPEEISKEKCRIAA--KLVGGAVMVEDTSLCFNA
Bot cin  26 DTIDL--QSQS------LDLVEIQG-TIEEISSDKCRRAA--EIIQGPVLVEDTCLCFNA
Neu cra  31 PAIQV--ENQA------LDLLEIQG-TLEEVTLDKCRRAAXXDLVQGPVLVEDTCLCFNA
Asp nig  26 TVIDV--ENQA------VDLPEIQG-TIEEIAREKCRRAA--EVVGGPVLTEDTALEFHA
Ust may  28 PNFPYELTNKD------LDLPEIQG-TTRDVAQAKCAAAAKXXALGGACITEDTALGFHA
Sac cer  25 QEVDNNNKTIHLINEA-LDLEELQDTDLNAIALAKGKQAVAALGKGKPVFVEDTALRFDE
Met jan  22 DLKDVEIEQIK------ISYPEIQG-TLEEVAEFGAKWVY--NILKKPVIVEDSGFFVEA
Bac sub  23 EPRGYDVKSLA--EIGFTEEIEETGHTFEENAILKAEAVAKXXAVNKMVIADDSGLSIDN
Dei rad  34 --APLGWQCEG---LGAVTLPEETGSTYEENAAXXLKACAAAMATGLPALADDSGIEVLA
Esc col  22 --SDFGLDIVAQTDLG-VDSAEETGLTFIENAILKARHAA--KVTGLPAIADDSGLAVDV
```

```
Gly max   81 LKGLPGPYIKWF--LQKLGHEGLNNLLMAY-----DDKSAYA---LCVFSFAA---GPDS
Pha vul  108 LKGLPGPYIKWF--LQKIGHEGLNNLLMAY-----DDKSAFA---LCVFSFAA---GPDS
Med tru   85 LKGLPGPYIKWF--LQKIGHEGLNNLLMAY-----DDKSAYA---LCVFSFAI---GPDS
Ara tha   84 LKGLPGPYIKWF--LEKLGHEGLNNLLMAY-----EDKSAYA---LCAFSFSR---GPGA
Sol tub   86 LKGLPGPYIKWF--LQKISHEGLNNLLMAY-----EDKTAYA---MCIFSLAL---GPNT
Sol lyc   87 LKGLPGPYIKWF--LQKIGHEGLNNLLMAY-----EDKTAYA---MCIFSLAL---GPNA
Tri aes   84 LKGLPGPYIKWF--LEKIGHEGLNNLLKAY-----EDKSAFA---MCIFSLAL---GPEE
Zea may   80 LKGLPGPYIKWF--LEKIGHEGLNNLLKAY-----EDKSAFA---MCIFSLAL---GPGE
Ory sat   81 LKGLPGPYIKWF--LEKTGHEGLNNLLLAY-----EDKSAFA---MCIFSL-----GPGE
Mar pol   82 LKGLPGPYVKWF--LQKTGHEGLNNLLAAY-----EDKSAYA---LCVFSLAL---GPDA
Sel moe   83 LHGLPG-FEKWF--LQKLGHEGLNNMLAAY-----KDKSAYA---LCVFSLAL---GPGF
Phy pat   82 LNGLPGPYVKWF--LMKTGHEGLNNLLAAY-----EDKTAYA---LCVFSLAL---GPDF
Dro mel   75 LEGLPGPYIKWF--LEKLQPEGLHRLLHGW-----ENKSAQA---ICTFGYCD---GVDA
Hom sap   80 LGGLPGPYIKWF--LEKLKPEGLHQLLAGF-----EDKSAYA---LCTFALST---GDPS
Sus scr   80 FGGLPGPYIKWF--LEKLKPEGLHQLLAGF-----QDKSAYA---LCTFALST---GDPN
Xen lae   79 LGGLPGPYIKWF--LEKIKPEGLHRMLEGF-----EDKSAIA---LCTFAYCN---GNPD
Dan rer   79 LEGLPGPYIKWF--LDKLKPEGLYKMLAGF-----EDKSAWA---LCTFAFCA---GKE-
Gal gal   79 LGGLPGPYIKWF--LEKLKPEGLYKLLAGF-----EDKSAYA---LCTFAFST---GNPE
Cae ele   73 MGGLPGPYIKWF--LKNLKPEGLHNMIAGF-----SDKTAYA---QCIFAYTE---GLG-
Vol car   79 LKGLPGPYIKWF--LEKLGHDGLNKMIAGF-----DDKTAYA---QCIFAYTT---GPEV
Chl rei   79 MHGLPGPYIKWF--LEKLGHDGLNRMIAGF-----EDKSAYA---QCIFAYTP---GPDT
Bot cin   75 LKELPGPYIKWF--MDALGHDGLNNMIAGF-----PDKSAQA---VCTFAYSE---GPGH
Neu cra   82 LKGLPGPYIKWF--MNSLGHEGLNNLLAAY-----EDKSAKA---VCTFGYSXXXAGPGH
Asp nig   75 LKGLPGPYIKSF--LDALGHEGLNKLLDSF-----ETRAAEA---VCTFAFSS---GPGS
Ust may   81 LGGLPGPYIKDF--MKTIGHDGLNKMLDGF-----EDRTASA---ICTFAYCAXXXGPDE
Sac cer   84 FNGLPGAYIKWF--LKSMGLEKIVKMLEPF-----ENKNAEA---VTTICFAD---SRGE
Met jan   73 LNGFPGTYSKFV--QETIGNEGILKLLEGK-----DNRNAYF---KTVIGYCD------E
Bac sub   81 LGGRPGVYSARYAGEQKDDQANINKVLSELKGIEKEQRTARFRXXXCALAVSI----PGE
Dei rad   89 LGGQPGVYSARF---GNVNSDVERNVLLLEKMRRHTDRXXXRAKFVSVLVLAY----PDG
Esc col   77 LGGAPGIYSARYSGEDATDLKNLQKLLETLKDVPDGQRQARF---HCVLVYLR--HAEDP

Gly max  128 EPI-TFSGKTPGKIVP-PRGPNDFG--WDPIFEP-DGYDQTYAQMPKEE---KNKIS---
Pha vul  155 EPI-TFSGKTPGKIVP-PRGPNDFG--WDPIFEP-EGYDQTYAQMPKEE---KNKIS---
Med tru  132 EPI-TFSGKTLGKIVP-PRGPNDFG--WDPVFQP-DGYDQTYAEMSKEE---KNKIS---
Ara tha  131 EPL-TFLGKTPGKIVP-ARGPTDFG--WDPVFQP-DGYDQTYAEMAKEE---KNKIS---
Sol tub  133 EPM-TFVGKTLGRIVP-ARGPNDFG--WDPIFQP-HGYDQTYAEMPKEE---KNKIS---
Sol lyc  134 EPM-TFVGKTQGRIVP-ARGPNDFG--WDPIFQP-HGYDQTYAEMPNEE---KHKIS---
Tri aes  131 EPI-TFVGKTAGKIVP-ARGPADFG--WDPVFQP-DGFEQTYAEMPKPE---KNQIS---
Zea may  127 EPI-TFVGKTAGKIVP-ARGPNYFG--WDPVFQP-DGFEQTYAEMPKSV---KNNIS---
Ory sat  126 EPM-TFVGKTAGKIVP-ARGPADFG--WDPVFQP-DGFDQTYAEMPKSV---KNQIS---
Mar pol  129 EPL-TFSGRTEGKIVP-ARGPGDFG--WDPVFQP-DGYDLTFAEMKKED---KNKIS---
Sel moe  129 EPI-TFVGRTEGKIVP-ARGPADFG--WDPVFQP-DGSDFTYAEMPKDE---KNKIS---
Phy pat  129 EPI-TFSGRTEGKIVP-ARGSGNFG--WDPIFQP-VGSDFTFAEMLKDE---KNKIS---
Dro mel  122 EPL-IFKGITEGVIVE-PRGPRDFG--WDPVFQP-SGYDKTYAELPKSE---KNTIS---
Hom sap  127 QPVRLFRGRTSGRIVA-PRGCQDFG--WDPCFQP-DGYEQTYAEMPKAE---KNAVS---
Sus scr  127 EPVRLFKGRTSGQIVV-PRGSRDFG--WDPCFQP-DGYEQTYAEMPKAE---KNTIS---
Xen lae  126 DTVLLFRGKTLGQIVL-PRGPRDFG--WDPCFQP-DGFQQTYAELPKEV---KNTIS---
Dan rer  125 EPVQLFRGITEGHIVE-PRGPRDFG--WDPCFQP-EGYDKTYAELPKEV---KNSIS---
Gal gal  126 EPVKLFKGQTHGVIVE-PRGPRDFG--WDPCFQP-DGYDQTYAELPKAV---KNSIS---
Cae ele  119 KPIHVFAGKCPGQIVA-PRGDTAFG--WDPCFQP-DGFKETFGEMDKDV---KNEIS---
Vol car  126 EPI-VFVGRTPGRIVA-ARGPPDFG--WDPIFEP-EGFETTYAEMDKET---KNKIS---
Chl rei  126 EPI-VFVGRTEGRIVQ-ARGPTDFG--WDPVFLP-DGFTDTYAEMDKTT---KNTIS---
Bot cin  122 EPI-IFQGRTDGKIVP-ARGPTAFG--WDPIFEY-EG--QTYAEMEKVE---KNKIS---
Neu cra  132 EPI-LFQGITDGKIVP-PRGPPNFG--WDAIFEY-EG--QTYAEMDKAE---KNKIS---
Asp nig  122 DPI-LFQGRTEGAIVR-PRGPANFG--WDPIFEH-QG--KTYAEMDKEE---KNRIS---
Ust may  131 QVH-LFEGRTEGVIVP-PRGPTHFG--WDPILEI-KGTGLTYAEMDPKQ---KNTLS---
Sac cer  131 YHF--FQGITRGKIVP-SRGPTTFG--WDSIFEPFDGHGLTYAEMSKDA---KNAIS---
Met jan  117 NGVRLFKGIVKGRVSE-EIRSKGYGFAYDSIFIP-EEEERTFAEMTTEE---KSQIS---
Bac sub  137 ETK-TVEGHVEGYIAEEPRGEYGFG--YDPIFIV-KDKDKTMAELTSDE---KNKISXXX
Dei rad  142 KLE-EYRGEVTGQLLEGPRGESGFG--YDPLFLP-DGSELSMGEMTLEQXXXKQAIS---
Esc col  132 TPL-VCHGSWPGVITREPAGTGGFG--YDPIFFV-PSEGKTAAELTREE---KSAIS---
```

```
Gly max  177 HRSKSLAL---VKSHFAEARF-----------------TFDVKNF----
Pha vul  204 HRSKSLAL---VKSHFAEAGY-----------------AFEINNV----
Med tru  181 HRSKSLAL---VKSHFAEAGY-----------------TFQI-------
Ara tha  180 HRYKSLAL---VKSHFKEAGY-----------------VFQTDDGTI--
Sol tub  182 HRGKALEL---VKLHFAEARY-----------------TFQTDSTTQN-
Sol lyc  183 HRGKALEL---VKLHFAEARY-----------------TFQTDSTT---
Tri aes  180 HRGRALAL---VKEHFASANY-----------------EVQGDGLA---
Zea may  176 HRGKALAL---VKEHFASASY-----------------TVQSNDST---
Ory sat  175 HRGKALAL---VKEHFAAANY-----------------KVQNDGSA---
Mar pol  178 HRRRALDK---VREHFYENDY-----------------APPV-------
Sel moe  178 HRRRALDK---VRDHFREYDF-----------------VVRNDDS----
Phy pat  178 HRRRALDK---VKEYFYDFNY-----------------APRSGNV----
Dro mel  171 HRYRALAL---LRQHFEKQDK-----------------LIN--------
Hom sap  177 HRFRALLE---LQEYFGSLAA----------------------------
Sus scr  177 HRFRALLE---LQEYFGSLTPPGAEAAHVISPLRPWSAVHSLSPNPVQ-
Xen lae  176 HRYRALKE---MSDYFIQNGT-----------------KV---------
Dan rer  175 HRYRALAA---LSEHFCQDNG-----------------APETKRSKHQD
Gal gal  176 HRYRALSE---LSAFFLQSNP-----------------TEAPSSPS---
Cae ele  169 HRAKALEL---LKEYFQNN------------------------------
Vol car  175 HRYRSLDL---LRSHLQHHSE-----------------KSS--------
Chl rei  175 HRYRSLDK---LRTYLLSHAA-----------------SK---------
Bot cin  169 HRFRALEK---LKTWLQEEA-----------------------------
Neu cra  179 HRAKALAKXXXLQEWFAKEMTA---------------------------
Asp nig  169 HRYKALVK---LQDWLADRRP----------------------------
Ust may  180 HRYKALTXXXLLQDYLVGLSKQN--------------------------
Sac cer  180 HRGKAFAQ---FKEYLYQNDF----------------------------
Met jan  169 HRKKAFEE---FKKFLLDRI-----------------------------
Bac sub  190 HRADALKK---LSKLLEA-------------------------------
Dei rad  195 HRGQALAA---LLAAHGA-------------------------------
Esc col  182 HRGQALKL---LLDALRNG------------------------------
```

**Figure S1.** Comparative sequence analysis of inosine triphosphate pyrophosphatases from several phylogenetically distant model organisms of all kingdoms of life.

Multiple alignment of sequences of inosine triphosphate pyrophosphatases (ITPA) from *Arabidopsis thaliana* and phylogenetically distant model organisms mentioned in the materials and methods. The alignment was generated using the MUSCLE algorithm in MEGAX. Shading indicating conserved residues was generated using BOXSHADE. Abbreviations and loci used for the phylogenetic analysis are as follows (Phytozome V12.1 locus identifiers for plant and algae sequences and NCBI locus identifiers for non-plant sequences are given): *Asp nig*, *Aspegillus niger* (XP_001398459.1); *Ara tha*, *Arabidopsis thaliana* (At4g13720); *Bac sub*, *Baccillus subitilis* (OIS57636.1); *Bot cin*, *Botrytis cinerea* (XP_024547817.1); *Cae ele*, *Caenorhabditis elegans* (NP_498121.1); *Chl rei*, *Chlamydomonas reinhardtii* (Cre02g095089.t1.1); *Dan rer*, *Danio rerio* (NP_001093456.1); *Dei rad*, *Deinococcus radiodurans* (WP_010886825.1); *Dro mel*, *Drosophila melanogaster* (NP_608890.1); *Esc col*, *Escherichia coli* (NAA10980.1); *Gal gal*, *Gallus gallus* (NP_001258859.1); *Gly max*, *Glycine max* (Glyma10G125100.1); *Hom sap*, *Homo sapiens* (AAK21848.1); *Mar pol*, *Marchantia polymopha* (Mapoly0026s0018.1); *Met jan*, *Methanocaldococcus jannaschii* (WP_064496428.1); *Med tru*, *Medicago truncatula* (Medtr1g047390.1); *Neu cra*, *Neurospora crassa* ( XP_955963.1); *Ory sat*, *Oryza sativa* (Os10g31940.1); *Phy pat*, *Physcomitrium patens* (Pp3c16_19640V3.1); *Pha vul*, *Phaseolus vulgaris* (Phvul.007G236950.1); *Sac cer*, *Saccharomyces cerevisiae* (AJR66682.1); *Sel moe*, *Selangiella moellendorffii* (e_gw1.0.2702.1); *Sol lyc*, *Solanum lycopersicum* (Solyc09g091420.2.1); *Sol tub*, *Solanum tuberosum* (PGSC0003DMT400076168); *Sus scr*, *Sus scrofa* (XP_020933236.1); *Tri aes*, *Triticum aestivum* (KAF6994464); *Ust may*, *Ustilago maydis* (XP_011387918.1); *Vol car*, *Volvox carteri* (Vocar0027s0182.1); *Xen lae*, *Xenopus laevis* (XP_018114098.1); *Zea may*, *Zea mays* (GRMZM2G020295_T02).

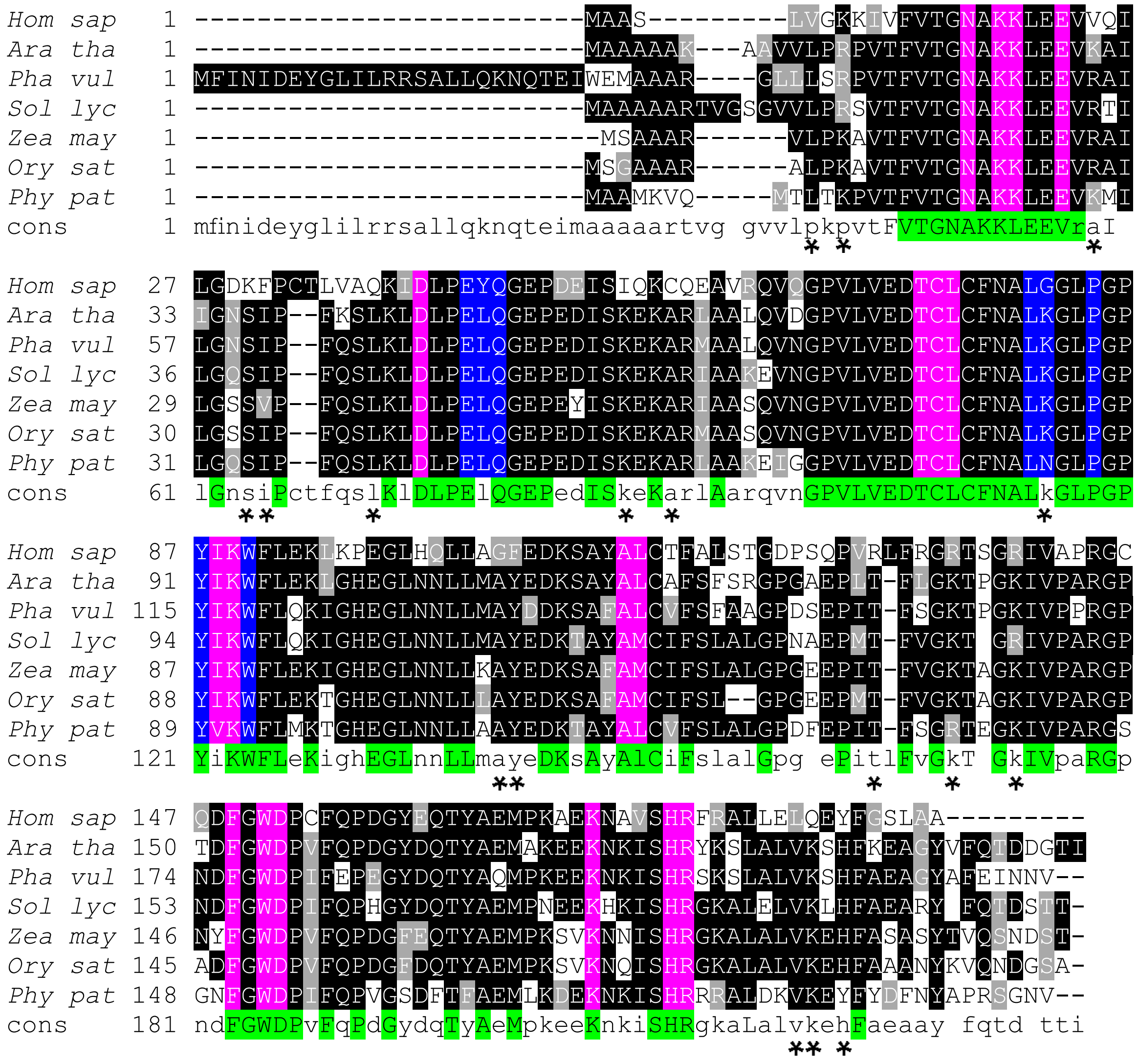


**Figure S2.** Comparative sequence analysis of inosine triphosphate pyrophosphatases.

Multiple alignment of sequences of inosine triphosphate pyrophosphatases (ITPA) from Homo sapiens and phylogenetically distant model plant organisms mentioned in the materials and methods. The alignment was generated using the MUSCLE algorithm in MEGAX. Shading indicating conserved residues was generated using BOXSHADE. Amino acids participating in substrate specificity and catalytic activity are indicated in pink, residues that are participating in enzyme dimerization are indicated in blue. Information about conserved amino acids is derived from Stenmark et al. (2006) and the Conserved Domain Database (Lu et al., 2020). Highly conserved amino acids occurring in all analyzed sequences are indicated in green in a consensus sequence below the analyzed sequences. Abbreviations and loci used for the phylogenetic analysis are as follows (Phytozome V12.1 locus identifiers for plant and algae sequences and NCBI locus identifiers for non-plant sequences are given): *Ara tha*, *Arabidopsis thaliana* (At4g13720); *Hom sap*, *Homo sapiens* (AAK21848.1); *Ory sat*, *Oryza sativa* (Os10g31940.1); *Phy pat*, *Physcomitrium patens* (Pp3c16_19640V3.1); *Pha vul*, *Phaseolus vulgaris* (Phvul.007G236950.1); *Sol lyc*, *Solanum lycopersicum* (Solyc09g091420.2.1); *Zea may*, *Zea mays* (GRMZM2G020295_T02). Some amino acid residues are specifically conserved in plant ITPAs, but not in the enzymes from other organisms and are indicated by an asterisk under the alignment (the given position refers to the ITPA sequence of A. thaliana): L12, R/K14, R/K30, S37, I/V38, L/V42, K58, A61, K85, A110, Y111, T134, K/R138, K142, V188, K/R189, H191.

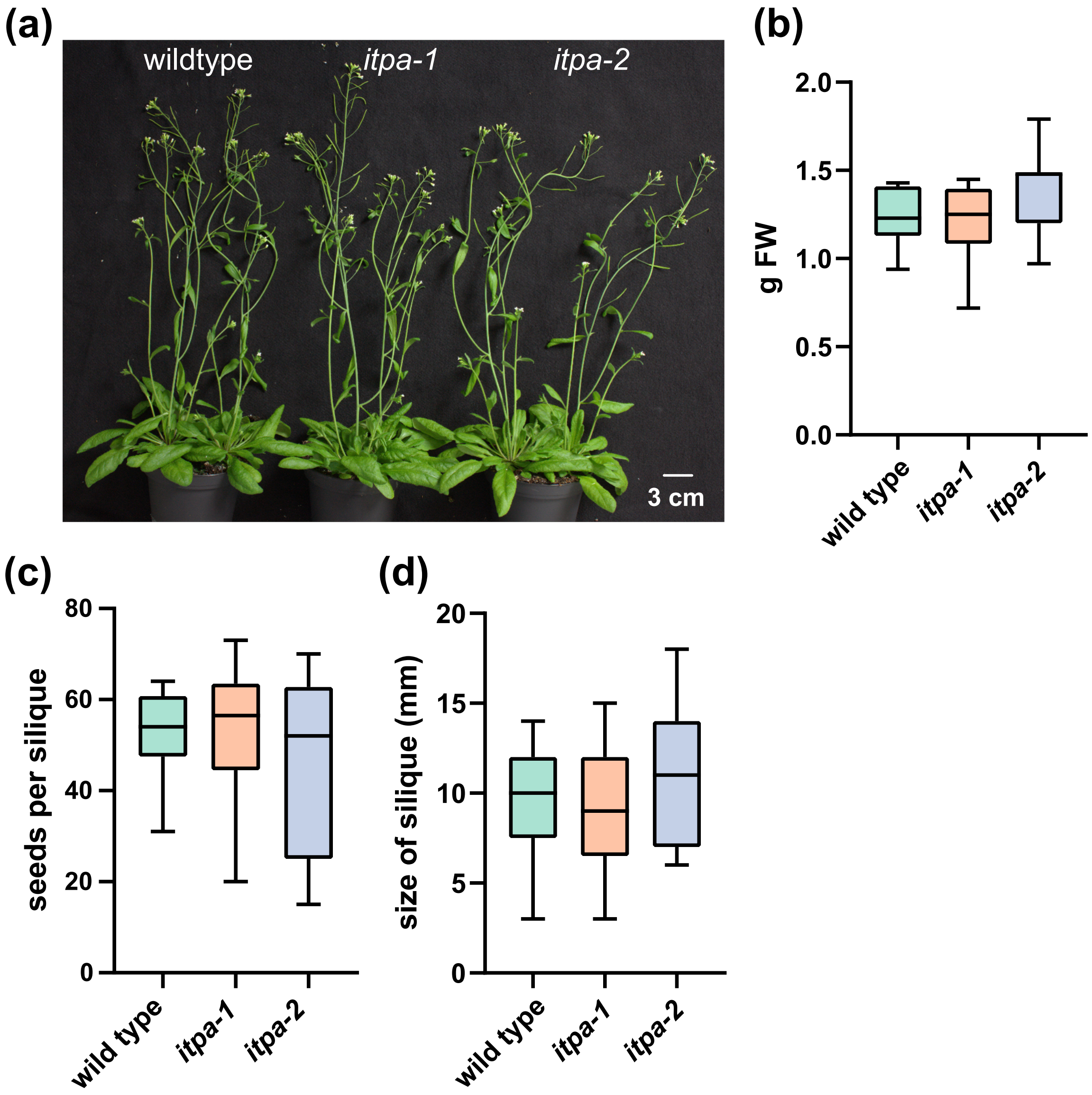


**Figure S3. Phenotyping of wild type and *itpa* mutant plants**

**(a)** Image of 35-day-old wild type, *itpa-1* and *itpa-2* plants
**(b)** Freshweight of 40-day-old wild type-, *itpa-1* and *itpa-2* plants. Rosettes and inflorescences were weighed without roots. Error bars are SD, $n$ = 9.
**(c)** Seeds per silique of wild type, *itpa-1* and *itpa-2* plants. Error bars are SD, $n$ = 12.
**(d)** Size of siliques of wild type, *itpa-1* and *itpa-2* plants. Error bars are SD, $n$ = 21.

Statistical analysis was performed by a two-sided Tukey's pairwise comparision using the sandwich variance estimator. Different letters indicate $p$ values < 0.05. All $p$ values can be found in Table S4.

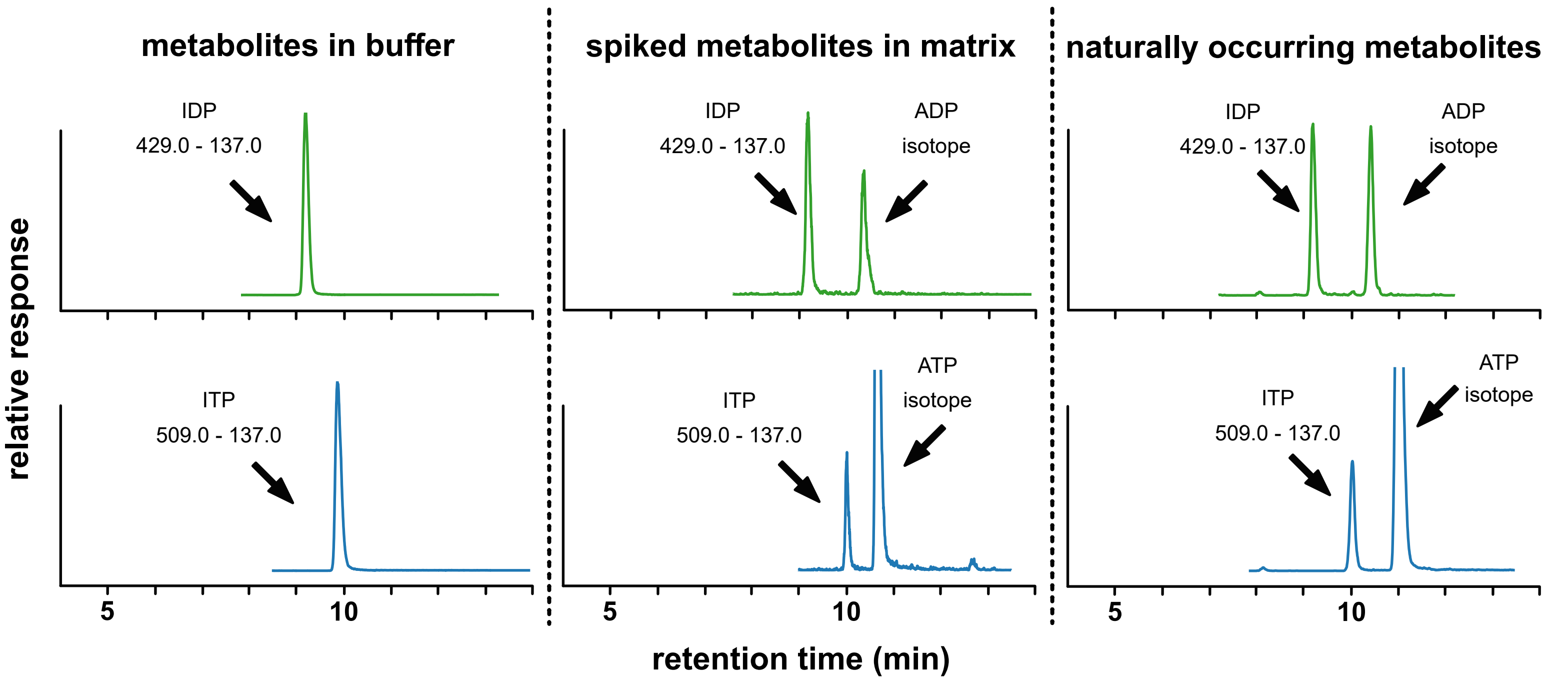


**Figure S4. Chromatograms of ITP and IDP analyzed by hypercarb chromatography.**

Representative depiction of Hypercarb multiple reaction monitoring (MRM) chromatograms of IDP (upper panel) and ITP (lower panel) in buffer (left column), wild type plant matrix isolated by SPE (middle column) or naturally occuring in *itpa* mutant plant matrix (right column). Chromatograms shown in green represent the IDP signal, while chromatograms shown in blue represent the ITP signal. MRM transitions used for quantification are given for the shown analytes. The signals of baseline separated ATP and ADP isotopes that are isobaric with ITP and IDP respectively are indicated by arrows.

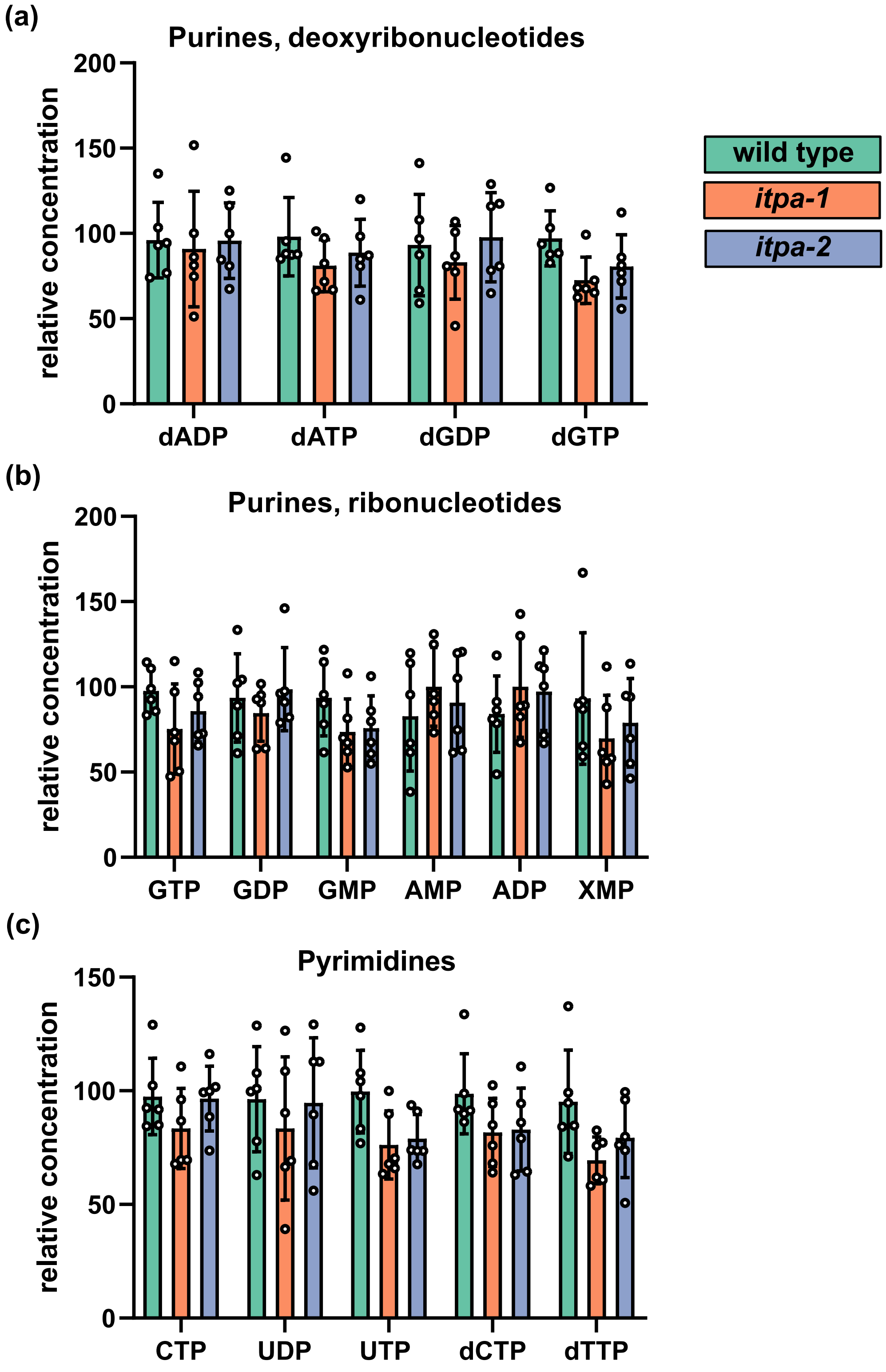


**Figure S5. Relative concentration of different nucleotides in rosette leaves of 35-day-old wild type and itpa mutant plants.**

Bars indicate the relative concentration of the respective metabolite in relation to the highest measured signal of the respective metabolite.

**(a)** Purine deoxynucleotides

**(b)** Purine ribonucleotides

**(c)** Pyridimiden nucleotides

Error bars are SD, *n* = 6, each replicate representing a different plant.

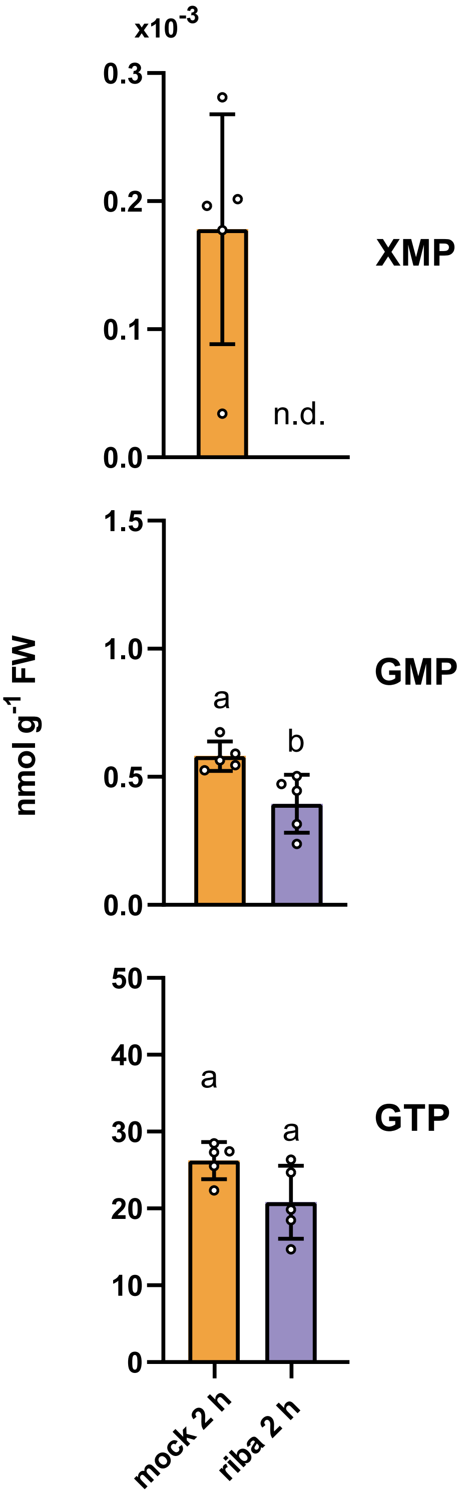


**Figure S6.** Manipulation of the IMP content in leaves of *N. benthamiana* alters the concentration of GMP and XMP, and in tendency also GTP.

XMP (upper panel), GMP (middel panel) and GTP (lower panel) in leaves of 21-day-old *N. benthamiana* plants.
The leaves were either mock infitrated with buffer on one leaf or infiltrated with buffer containing ribavirin (500 µM) on the opposite leaf. Error bars are SD, $n = 5$, every replicate representing a leaf from an independent plant. Statistical analysis was performed by a two-sided Tukey's pairwise comparision using the sandwich variance estimator. Different letters indicate $p$ values < 0.05. All $p$ values can be found in the Supplemental Table 1. n.d. = not detected.

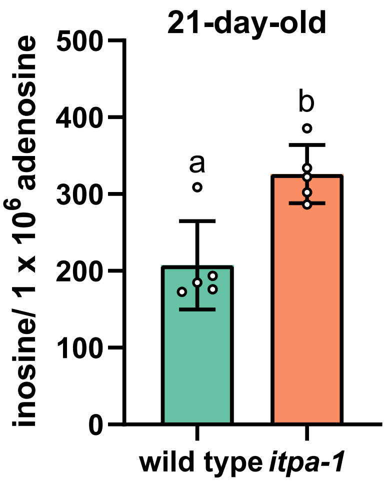


**Figure S7. Concentration of inosine in isolated RNA of wild type and *itpa-1* plants.**

Molecules of inosine per million adenosine molecules in RNA isolated from 21-day-old arabidopsis rosette leaves. Error bars are SD, *n* = 5, every replicate representing a different plants. Statistical analysis was performed by a two-sided Tukey's pairwise comparision using the sandwich variance estimator. Different letters indicate *p* values < 0.05. All *p* values can be found in the Table S4.

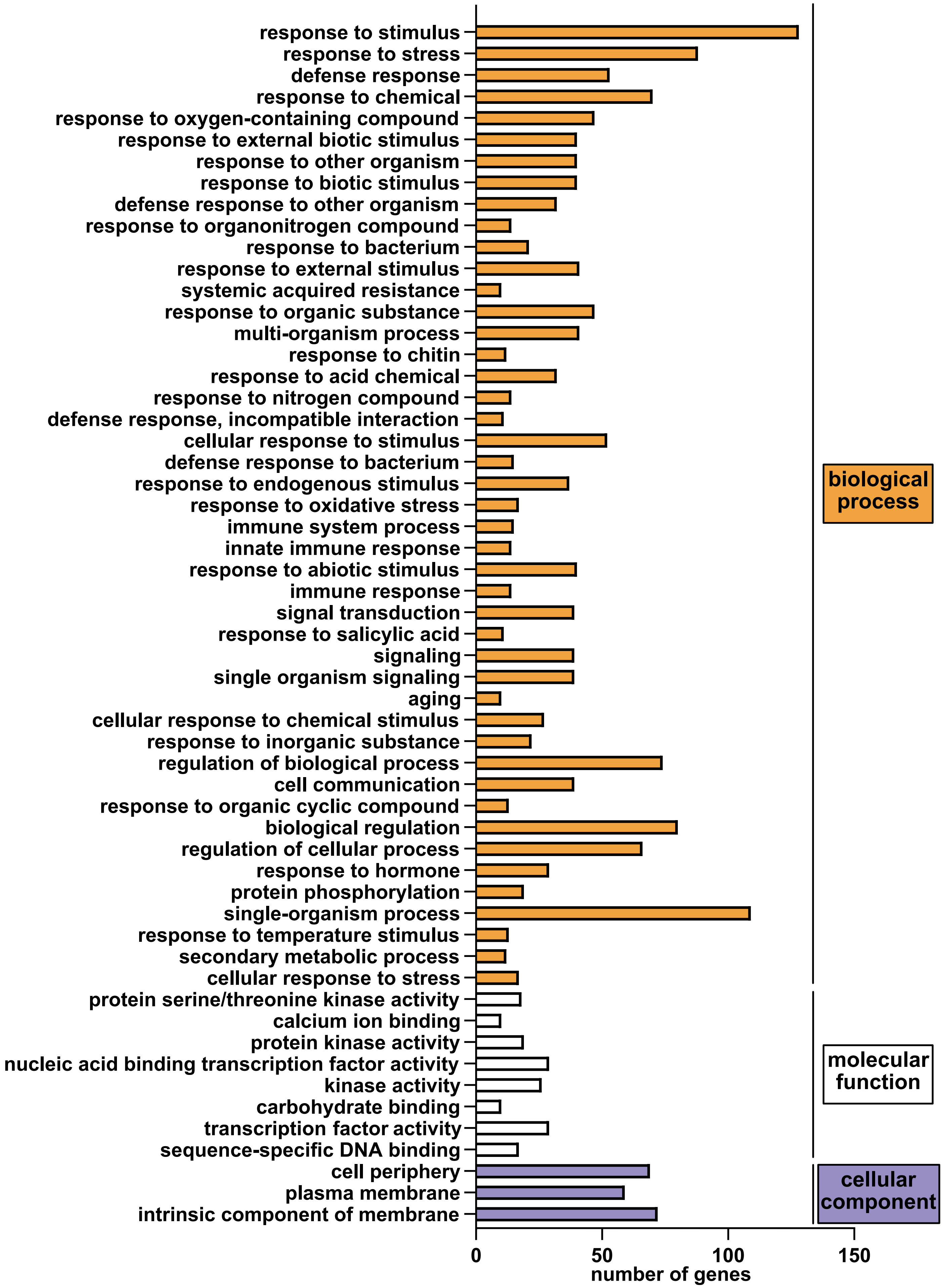


**Figure S8. Loss-of-function of ITPA leads to differential upregulation of transcripts associated with GO:terms of biological processes, molecular functions and cellular components.**

RNA was isolated from 35-day-old Arabidopsis rosettes. GO:term analysis of differentially upregulated genes ($log_2$ FC > 2; FDR < 0.05) in *itpa-1* plants compared to wildtype plants. The analysis was performed with AgriGO v2.0 based on singular enrichment analysis. Statistics were performed using hypergeometric test after Hochberg with significant level adjusted *P* < 0.01. Only GO:terms with a minimum of 10 mapping entries are displayed.
