## Supplementary material for "An inosine triphosphate pyrophosphatase safeguards plant nucleic acids from aberrant purine nucleotides": Table S1,5,6

**Supplemental Table 1**: List of primers used in this study

| **Name** | **Sequence** |
| --- | --- |
| **Primers to verify genetic status and transcript abundance** | |
| At4g13720_tDNA_diag_fwd (P-1628) | GGAACACATAGCCAGCCTCCTT |
| At4g13720_tDNA_diag_rvs (P-1629) | GGACTGATTTGTAGAAGTGGCC |
| LBb1.3 (P-1316) | ATTTTGCCGATTTCGGAAC |
| At4g13720_transcript_detection_fwd (P-2487) | CGTTTGTGACTGGAAATG |
| At4g13720_transcript_detection_rvs (P-2488) | GCTAAAGACTTGTACCTATGTG |
| control actin2fwd (1033) | GTGAACGATTCCTGGACCTGC |
| control actin2frvs (1034) | GAGAGGTTACATGTTCACCACAAC |
| **Primers for the assembly of C-terminal strep-tag and YFP fusion proteins** | |
| At4g13720_sense (P-1075) | agaGAATTCAAAATGGCGGCGGCGGCGGCGAAA |
| At4g13720_anti (P-1076) | cgaCCCGGGAATGGTACCATCATCTGTCT |
| **Primers for the assembly of CRISPR-Cas vector** | |
| At4g13720_gR1_f (P-1657) | attgTGAGCTCCAAGGTGAGCCTG |
| At4g13720_gR1_r (P-1658) | aaacCAGGCTCACCTTGGAGCTCA |
| At4g13720_gR2_f (P-1659) | attgCTAGAGAAGCTTGGTCACGA |
| At4g13720_gR2_r (P-1660) | aaacTCGTGACCAAGCTTCTCTAG |
| gRNA_MAR1_At_Cas9_fwd (P-1436) | ATTGgataattgctggaggccctg |
| gRNA_MAR1_At_Cas9_rev (P-1437) | AAACcagggcctccagcaattatc |
| At5g26820_gR2_f (P-1665) | attgGCCACTCCTGCTCACCCTGA |
| At5g26820_gR2_r (P-1666) | aaacTCAGGGTGAGCAGGAGTGGC |
| At4g13720_capseq_fwd1 (P-1774) | GTAAAACGACGGCCAGTGCGTACGGCGTACCC |
| At4g13720_capseq_rev1 (P-1775) | GCAACAGTACATAAGAAGCAAGC |
| At4g13720_capseq_fwd2 (P-1776) | GTAAAACGACGGCCAGTGCCTAGGACTAACTGTATTGGAGC |
| At4g13720_capseq_rev2 (P-1777) | CCATCAGTAAGTTGTTCAGACCTG |
| MoClo-CRISPR-control-A5 fw.(P-1164) | AAAGCTGCAAATGTTACTGA |
| MoClo-CRISPR-control-A5 rev (P-1165) | GGCAACCTCGCATGAAAATAGTA |

**Supplemental Table 5**: Substrates tested for biochemical activity with ITPA

| **Substrate (200 µM)** | **Activity** |
| --- | --- |
| dITP | yes |
| ITP | yes |
| XTP | yes |
| IDP | yes |
| dADP | yes |
| dGDP | yes |
| dCDP | no |
| dATP | no |
| dGTP | no |
| dCTP | no |
| dTTP | no |
| dUTP | no |
| ATP | no |
| GTP | no |
| UTP | no |
| CTP | no |
| NAD | no |
| NADH | no |
| NADP | no |
| pNPP | no |

**Supplemental Table 6**: Ratios of (deoxy)inosine per million molecules (deoxy)adenosine in DNA and RNA, respectively

|  | **deoxyinosine/ 10^6^ deoxyadenosine** | | |
| --- | --- | --- | --- |
|  | **wild type** | ***itpa-1*** | ***itpa-2*** |
| **DNA (7-days-old)** | 290 ± 96 | 699 ± 223 | 756 ± 268 |
| **DNA (56-days-old)** | - | 837 ± 192 | 1534 ± 661 |
|  | **inosine/ 10^6^ adenosine** | | |
|  | **wild type** | ***itpa-1*** | ***itpa-2*** |
| **RNA (35-days-old)** | 404.6 ± 117.4 | 660 ± 85 | 909 ± 240 |
