## Supporting methods for "An inosine triphosphate pyrophosphatase safeguards plant nucleic acids from aberrant purine nucleotides"

**Genotyping and cloning**

*Arabidopsis* T-DNA insertion mutants (*itpa-1*) and respective wild type plants were identified by PCR using the following primer combinations: P1628 and P1629 (wild type reaction); P1628 and P1316 (T-DNA reaction). The PCR product of the T-DNA reaction was sent for sequencing. The T-DNA insertion was found to be positioned as described on the Signal website (<http://signal.salk.edu/cgi-bin/tdnaexpress>). Gene-specific mRNA level were determined by extracting total RNAs from seedlings of wild type and knockout plants and subsequent reverse transcription (Revert Aid H Minus first strand cDNA synthesis kit, ThermoFisher). PCRs were facilitated using T-DNA insertion flanking primers P2487 and P2488. An amplification of *actin2* was used as control with the primers 1033 and 1034. The PCR was carried out with 35 cycles using an annealing temperature of 55°C. Vectors for CRISPR-induced editing of the ITPA encoding locus At4g13720 were constructed as previously described (Rinne et al., 2021). We used two gRNAs (target sequences: TGAGCTCCAAGGTGAGCCTG; CTAGAGAAGCTTGGTCACGA), each expressed by an AtU6-26-promoter on the same vector. The final vector contained (I) Selectable phosphinotricin resistance cassette encoding a phosphinotricin acetyltransferase; (II) Cas9 under the controle of the EC1.2-promoter for egg cell-specific Cas9-expression (Wang et al., 2015); (III) the gRNA-expression cassette and (IV) green fluorescent protein (GFP) expressed by seed specific At2S3-promoter (Aliaga-Franco et al., 2019) for selection of non-glowing T-DNA-free seeds. The vector was transformed into *Agrobacterium tumefaciens* strain AGL-1. *Arabidopsis thaliana* Col-0 plants were grown as described above until flowers developed and transformed by floral dip (Clough and Bent, 1998). T1 seeds were harvested and screened for transgenicity by spraying 1:1000 diluted phosphinotricin (BASTA®, Bayer). Resistant plants were grown until seeds developed and T2-seedlings were genotyped for mutations in the ITPA locus. Genotyping was performed via amplified fragment length polymorphism (AFLP) analysis of the genomic area around the targeted sites as described previously (Rinne et al., 2021). The first target site located in the second exon was amplified with P1774 and P1775 and the second target site located in the fifth exon was amplified with P1776 and P1777. Mutations were confirmed by Sanger sequencing (Microsynth-Seqlab). Plants harboring mutations in the ITPA locus were grown further until seeds developed. To obtain T-DNA-free T3 seeds, non-glowing seeds were selected and germinated. Absence of the T-DNA was confirmed by PCR-amplification with P1164 and P1165 and inheritance of mutations in the ITPA locus were confirmed by Sanger sequencing. Plants that showed no DNA-amplification with T-DNA-specific oligonucleotides were used for further experiments.

The ITPA coding region was amplified from cDNA using the primers P1075 and P1076. The fragment was cloned into pXCScpmv-HA-Strep (V69, Claus-Peter Witte et al., 2004) for protein production and into pXCS-YFP (V36, Myrach et al., 2017; Dahncke and Witte, 2013) for localization studies.

**Enzymatic assays**

The enzymatic activity of ITPA was measured at 360 nm using an UV-2700 UV-Vis-spectrophotometer (Shimadzu). Resulting data were analyzed with Graph Pad Prism 4, determining kinetic constants fitting the data to the Michaelis-Menten equation. Specific activities were screened with 200 μM of the respective substrate, enzymatic constants for ITP were determined using concentrations of 1.5, 3.0, 6.25, 12.5, 25.0 and 50.0 μM ITP; for dITP concentrations of 1, 3, 6, 15, and 30 μM dITP were used.

AMPK3 and 4 activity was determined using a coupled enzyme assay with pyruvate kinase and lactate dehydrogenase (LDH, Sigma, P0294), measuring the production of NAD^+^ that results in absorption-decrease at 340 nm (AMPK3) or 1 μl purified enzyme (AMPK4) were used in a total volume of 300 μl containing 40 mM Tris-SO_4_, 20 mM KCL, 4 mM MgCl_2_, 2 mM phosphoenolpyruvate, 0.17 mM NADH, 2 μl 1:50 diluted coupling enzyme solution (600-1000 U/ml PK and 900-1400 U/ml LDH). The reaction was started by adding ATP to a final concentration of 1 mM and measured by a microplate spectrophotometer (Multiskan GO, Thermo Scientific) at 22°C for 20 min. The absorption change was calculated from the linear phase and subtracted by the change in a blank-sample, containing everything but the substrate. Absolute amounts were calculated utilizing the Beer-Lambert law.

**Quantitative determination of (deoxy)inosine per (deoxy)adenosine level in total RNA and DNA**

Briefly, 1 g plant material was frozen in liquid nitrogen and ground in a mortar with 50 mg polyvinylpolypyrrolidone. 1.5 ml CTAB Buffer (2% CTAB; 20 mM EDTA; 1.4 M NaCl; 150 mM Tris-HCL pH 7.5) were added and the material was thawed with occasional gentle stirring, transferred to 2 x 2 ml reaction tubes and placed in a water bath for 10 min at 60°C. 0.5 ml chloroform:isoamylacohol (24:1) was then added and the tubes were inverted 10 times. The tubes were centrifuged at room temperature for 5 min at 21000 x *g* and the supernatant was transferred to a new 2 ml tube containing 0.5 ml phenol:chloroform:isoamylacohol (25:24:1). This solution was inverted thoroughly and centrifuged as stated before. The phenol-step was repeated and the resulting supernatant was transferred to a tube containing 0.5 ml chloroform, inverted and centrifuged for 3 min at room temperature at 21000 x *g*. The supernatant was then pooled in a 2 ml reaction tube and 0.6 times the volume of isopropanol was added and centrifuged at 21000 x *g* for 2 min to precipitate the nucleic acids. The supernatant was discarded and the pellet was resolved in 600 μl TE buffer (5 mM, pH 7.5) containing 50 μg/ml RNAse A and incubated at 60 °C for 10 min in a water bath. Precipitates were removed by centrifugation at 21.000 x *g* for 3 min. 60 μl 3 M sodium acetate (pH 5.2) and 1.2 ml ethanol were added to the supernatant and the supernatant was spun down at 21000 x *g* for 5 min to precipitate the DNA. The pellet was washed with 70% ethanol and resuspended in 100 μl Tris buffer (5 mM). Nucleic acids concentrations were determined using a spectrophotometer (Nanophotometer, Implen).

A Poroshell 120 SB-C18 (2.7 μm, 2.1 × 150 mm; Agilent Technologies) was employed for chromatographic separation using a gradient employing solvent A (0.0075% formic acid in water) and solvent B (0.0075% formic acid in methanol) (0-3min min, 0% solvent B; 3.0-8.0 min 11% solvent B; 8.0-8.25 min, 50% solvent B; 8.25-10.25 min, 50% solvent B; 10.25-10.50 min, 0% solvent B until 13.5 min). A flowrate of 0.35 ml/min was applied and 10 μl digested RNA were injected. The source parameters were set as following: Positive mode, gas temperature 80°C, gas flow 13 L/min, nebulizer 30 psi, sheath gas temperature 275°C, sheath gas flow 11 L/min, capillary voltage 2500 V and nozzle voltage 500 V.

**Sequence Analysis and Phylogenetic Analysis**

Alignments were generated using the Muscle algorithm in MEGAX. The alignment was used to construct a maximum likelihood tree using MEGAX (Kumar et al., 2018). Initial tree(s) were obtained automatically by applying Neighbor-Join and BioNJ algorithms to a matrix of pairwise distances estimated using a JTT model, and then selecting the topology with superior log likelihood value (-5225.0). All positions with less than 95% coverage were eliminated, resulting in 173 positions in the final dataset. A discrete Gamma distribution was used to model evolutionary rate differences among sites (5 categories (+G, parameter = 0.9102)). Numbers at branches indicate the percentage of trees in which associated taxa clustered together and were calculated by a bootstrap analysis (1000 bootstraps). The alignment used to construct the phylogenetic tree can be found in Fig. S1. Conserved residue shading was performed using Boxshade (http://www.ch.embnet.org/software/BOX_form.html). Conserved domains were identified using information from crystallography studies of the human ITPA (Stenmark et al., 2007), as well as bioinformatics analysis using the Conserved Domain Database (Lu et al., 2020). Abbreviations and loci used for the phylogenetic analysis are as follows (Phytozome V12.1 locus identifiers for plant and algae sequences and NCBI locus identifiers for non-plant sequences are given): *Asp nig*, *Aspegillus niger* (XP_001398459.1); *Ara tha*, *Arabidopsis thaliana* (At4g13720); *Bac sub*, *Baccillus subitilis* (OIS57636.1); *Bot cin*, *Botrytis cinerea* (XP_024547817.1); *Cae ele*, *Caenorhabditis elegans* (NP_498121.1); *Chl rei*, *Chlamydomonas reinhardtii* (Cre02g095089.t1.1); *Dan rer*, *Danio rerio* (NP_001093456.1); *Dei rad*, *Deinococcus radiodurans* (WP_010886825.1); *Dro mel*, *Drosophila melanogaster* (NP_608890.1); *Esc col*, *Escherichia coli* (NAA10980.1); *Gal gal*, *Gallus gallus* (NP_001258859.1); *Gly max*, *Glycine max* (Glyma10G125100.1); *Hom sap*, *Homo sapiens* (AAK21848.1); *Mar pol*, *Marchantia polymopha* (Mapoly0026s0018.1); *Met jan*, *Methanocaldococcus jannaschii* (WP_064496428.1); *Med tru*, *Medicago truncatula* (Medtr1g047390.1); *Neu cra*, *Neurospora crassa* (XP_955963.1); *Ory sat*, *Oryza sativa* (Os10g31940.1); *Phy pat*, *Physcomitrium patens* (Pp3c16_19640V3.1); *Pha vul*, *Phaseolus vulgaris* (Phvul.007G236950.1); *Sac cer*, *Saccharomyces cerevisiae* (AJR66682.1); *Sel moe*, *Selangiella moellendorffii* (e_gw1.0.2702.1); *Sol lyc*, *Solanum lycopersicum* (Solyc09g091420.2.1); *Sol tub*, *Solanum tuberosum* (PGSC0003DMT400076168); *Sus scr*, *Sus scrofa* (XP_020933236.1); *Tri aes*, *Triticum aestivum* (KAF6994464); *Ust may*, *Ustilago maydis* (XP_011387918.1); *Vol car*, *Volvox carteri* (Vocar0027s0182.1); *Xen lae*, *Xenopus laevis* (XP_018114098.1); *Zea may*, *Zea mays* (GRMZM2G020295_T02).

**SUPPORTING METHODS REFERENCES**

**Aliaga-Franco, N., Zhang, C., Presa, S., Srivastava, A.K., Granell, A., Alabadí, D., Sadanandom, A., Blázquez, M.A., and Minguet, E.G.** (2019). Identification of transgene-free CRISPR-edited plants of rice, tomato, and arabidopsis by monitoring DsRED fluorescence in dry seeds. Frontiers in Plant Science **10**: 1150.Floral dip: a simplified method for Agrobacterium-mediated transformation of Arabidopsis thaliana. The Plant Journal for Cell and Molecular Biology **16** (6): 735–743.

**Clough, S.J., and Bent, A.F.** (1998). Floral dip: a simplified method for Agrobacterium-mediated transformation of Arabidopsis thaliana. The Plant Journal for Cell and Molecular Biology **16** (6): 735–743.

**Kumar, S., Stecher, G., Li, M., Knyaz, C., and Tamura, K.** (2018). MEGA X: Molecular evolutionary genetics analysis across computing platforms. Molecular Biology and Evolution **35** (6): 1547–1549.

**Lu, S., Wang, J., Chitsaz, F., Derbyshire, M.K., Geer, R.C., Gonzales, N.R., Gwadz, M., Hurwitz, D.I., Marchler, G.H., Song, J.S., Thanki, N., Yamashita, R.A., Yang, M., Zhang, D., Zheng, C., Lanczycki, C.J., and Marchler-Bauer, A.** (2020). CDD/SPARCLE: the conserved domain database in 2020. Nucleic Acids Research **48** (D1): D265-D268.

**Rinne, J., Witte, C.-P., and Herde, M.** (2021). Loss of MAR1 Function is a Marker for Co-Selection of CRISPR-Induced Mutations in Plants. Frontiers in Genome Editing **3**: 723384.

**Stenmark, P., Kursula, P., Flodin, S., Gräslund, S., Landry, R., Nordlund, P., and Schüler, H.** (2007). Crystal structure of human inosine triphosphatase. Substrate binding and implication of the inosine triphosphatase deficiency mutation P32T. Journal of Biological Chemistry **282** (5): 3182–3187.

**Wang, Z.-P., Xing, H.-L., Dong, L., Zhang, H.-Y., Han, C.-Y., Wang, X.-C., and Chen, Q.-J.** (2015). Egg cell-specific promoter-controlled CRISPR/Cas9 efficiently generates homozygous mutants for multiple target genes in Arabidopsis in a single generation. Genome Biology **16**: 144.
